# Integrated K^+^ channel and K^+^-Cl^−^ cotransporter functions regulate fin proportionality in zebrafish

**DOI:** 10.1101/621243

**Authors:** Jennifer S. Lanni, David Peal, Laura Ekstrom, Haining Chen, Caroline Stanclift, Margot Bowen, Adriana Mercado, Gerardo Gamba, Kristopher T. Kahle, Matthew P. Harris

## Abstract

The coordination of growth during development establishes proportionality within and among the different anatomic structures of organisms. Innate memory of this proportionality is preserved, as shown in the ability of regenerating structures to return to their original size. Although the regulation of this coordination is incompletely understood, mutant analyses of zebrafish with long-finned phenotypes have uncovered important roles for bioelectric signaling in modulating growth and size of the fins and barbs. To date, long-finned mutants identified are caused by hypermorphic mutations, leaving unresolved whether such signaling is required for normal development. We isolated a new zebrafish mutant, *schleier*, with proportional overgrowth phenotypes caused by a missense mutation and loss of function in the K^+^-Cl^−^ cotransporter Kcc4a. Genetic depletion of Kcc4a in wild-type fish leads to a dose-dependent loss of growth restriction in fins and barbs, supporting a requirement for Kcc4a in regulation of proportion. Epistasis experiments suggest that Kcc4a and the two-pore potassium channel Kcnk5b both contribute to a common bioelectrical signaling response in the fin. These data suggest that an integrated bioelectric signaling pathway is required for the coordination of size and proportion during development.

**Graphical Abstract:** 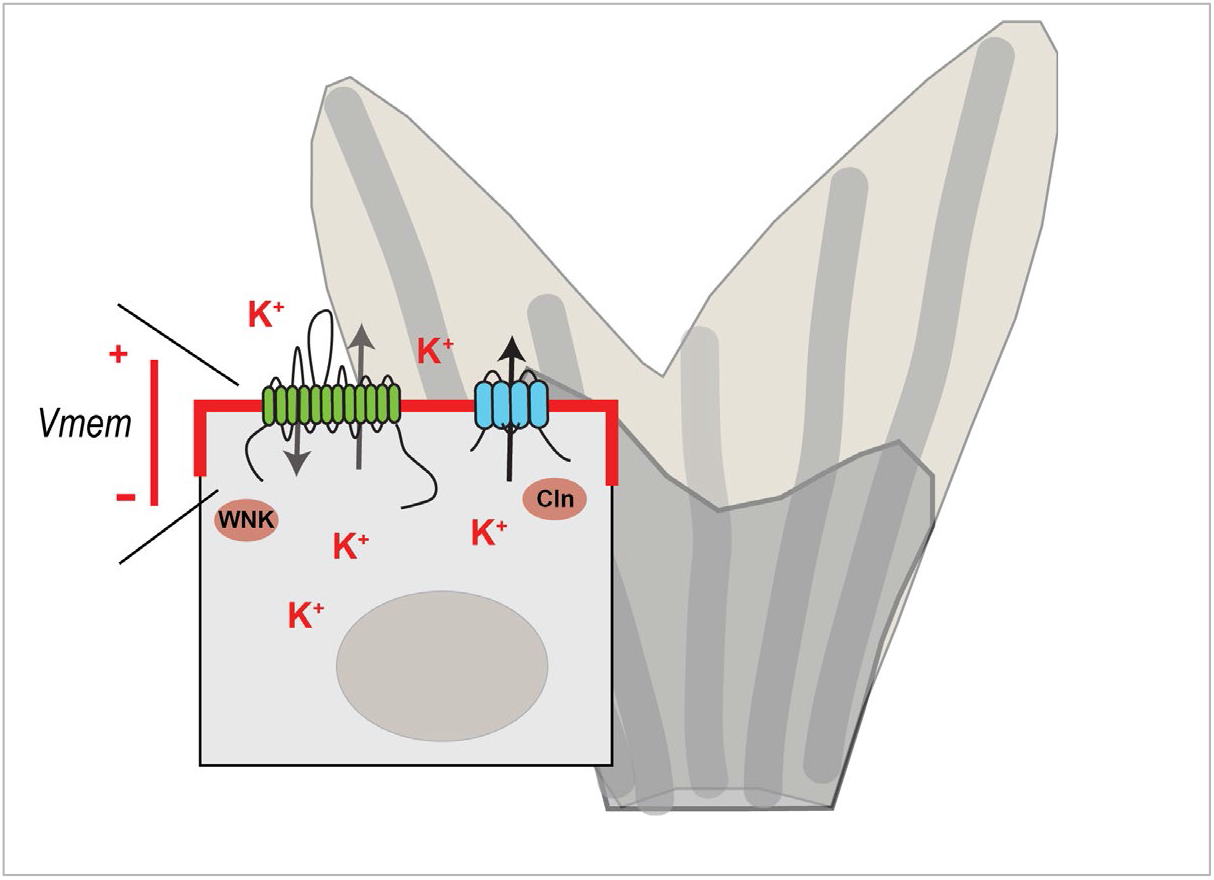

## Introduction

Anatomical systems depend on scaling properties to function such that as size increases, structure of tissues and their interactions change to accommodate (Huxley and Teissier, 1936; Huxley, 1932). In many systems this scaling is a property of morphogenetic fields and differential read out as size changes; in other contexts, scaling has been attributed to space filling models and reaction diffusion-based patterning (Umulis and Othmer, 2013; Othmer and Pate, 1980). A lasting question in scaling is how overall size is determined such that there is a relative growth of structures in relation to total body size (Vollmer, et al., 2017) - in other words, how do structures know when to stop growing? These relative proportions are established in development and maintained such that they are renewed in regenerating systems. When tissues fail to coordinate these patterning functions within structures or have dysregulated overall growth, aberrant cell behavior and neoplasm often result.

Genetic analyses have been utilized to great effect over the last several decades to identify the molecules that regulate size and growth of structures during development and regeneration. Many of these genetic factors act as components of transcriptional networks functioning downstream of intercellular signaling peptides and hormones that control systemic regulation of local tissue and cell behavior. In growth regulation, genes such as *hippo/yap* and insulin growth factors have been tied to overgrowth and size regulation (reviewed in (Watt, et al., 2017; Gokhale and Shingleton, 2015). However, the overarching mechanism of developmental signaling that mediates coordinated growth and relative proportion remains enigmatic.

Recently, research has revealed the importance of additional pathways such as bioelectric signaling in governing growth and coordinated development within and between tissues. Bioelectric signals stemming from local cell and tissue sources as well as broader systemic sources such as electrical fields have been found to be associated with organized biological structures such as tissues, organs and organisms (Levin, et al., 2017). At the cellular level, bioelectric signaling can be observed to regulate the proliferation rate of cells via alterations in transmembrane voltage gradients (V_mem_). These changes in V_mem_ can also be transduced within tissue layers by cellular junctions to create tissue-wide bioelectric gradients that effect changes in large-scale patterning (Levin, et al., 2017). Despite this progress, much remains to be elucidated about bioelectrical signaling networks, from the specific molecules that determine transmembrane voltage gradients to the mechanisms by which these gradients are expressed in body patterning and growth. Further, the informative nature of such bioelectric signals in development has been an open question for over the last 120 years of research, and it remains unclear if these signals are a consequence of, or a specific regulator of development and tissue/organ homeostasis.

Forward mutagenesis screens in zebrafish have identified several genes that are implicated in both bioelectric signaling and body patterning. The distinctive striped pigment pattern in adult zebrafish is altered by mutation of the potassium channel *kcnj13 (obelix/jaguar,* (Iwashita, et al., 2006)); this phenotype has been shown to be due to the absence of cell depolarization normally triggered by contact between pigment cells (Inaba, et al., 2012). Similarly, mutations in the gap junction proteins *connexin 39.4 (luchs)* and *connexin 41.8 (leopard)* also affect pigment stripe width or integrity (Irion, et al., 2014; Watanabe, et al., 2006). Interestingly, adult fin size is also regulated by proteins with conductive properties. Mutation of the *connexin 43* gene causes a short fin phenotype (Iovine, et al., 2005) that is proportional to loss of junctional conductance (Misu, et al., 2016; Hoptak-Solga, et al., 2007). We have previously identified the two-pore potassium channel *kcnk5b* as an important mediator of fin size regulation in zebrafish. Specifically, fish with activating mutations in *kcnk5b* exhibit proportional overgrowth of the fins and barbels that is wholly dependent upon channel conductivity. *Kcnk5b* expression is also required for fin overgrowth induced by inhibition of calcineurin, providing a link between bioelectric signaling and transcriptional regulation by calcineurin. The cases of enhanced growth or patterning seen in the long-finned or broad-stripe phenotypes described above are quite informative, as loss of structures can occur for many genetic and physiological reasons but enhanced growth indicates specificity of regulation. However, the chemical treatments and genetic alleles identified thus far that cause proportional fin overgrowth are all gain-of-function and may thereby reflect a supra-physiological response of tissues to novel growth enhanced conditions, whether bioelectrical or chemical.

In this work, we identify a novel zebrafish mutant, *schleier*, that affects both fin length and pigment pattern in the adult fish and is similar in fin phenotype to gain of function mutants of *kcnk5b* (Perathoner, et al., 2014). We show that these phenotypes are due to inactivation of the potassium-chloride cotransporter *slc12a7/kcc4a*, which carries out electroneutral symport across the plasma membrane. While *slc12a7* paralogues have not previously been studied in zebrafish, the orthologue of *slc12a7a* has a broad expression pattern in mammals and is linked to diverse functions in the kidney, inner ear, and gastric and prostate glands (Boettger, et al., 2002) (Marcoux, et al., 2017). Importantly, we demonstrate that proportional growth of zebrafish appendages encompassing fins and barbels is dependent on *slc12a7a*/*kcc4a*; thus, potassium regulation is essential for establishing the normal size of these appendages in development. Further, we find that the enhanced growth effects stemming from loss of Kcc4 function are attenuated by Kcnk5b inactivation, suggesting that these two channels act in a similar pathway. A dominant negative mutation in Kcc4a also disrupts normal pigment pattern in the adult fish and resembles zebrafish mutants affecting gap junction function. We also show that Kcc4 inactivation in fins results in a dramatic change in fin vasculature, providing a potential mechanistic connection between vascularization, potassium regulation and fin growth.

Our work highlights the growing importance of bioelectric signaling in growth regulation and coordinated patterning of complex structures. Through our systematic mutagenesis analysis in the zebrafish, we have identified a genetic signaling cascade combining environmental, systemic, and canonical intracellular signaling networks through integration of bioelectric signaling in development.

## Results

### Identification and characterization of *schleier* mutant zebrafish

In a screen to identify dominant modifiers of *longfin^dt2^*(*lof*) mutant zebrafish, a mutant was identified that showed additive effects with *lof* on fin length (**Supplementary Figure 1**). This mutation segregated away from the *lof* locus and was isolated. Heterozygous mutants exhibit changes in proportion of all fins and barbels (**Figure 1a, 1e**). This effect is dose dependent with homozygotes exhibiting slightly dysmorphic fins due to internal fractures of the fin rays. (**Figure 1c**). This mutant closely resembles fin overgrowth morphologies observed in gain-of-function mutations in the potassium channel *kcnk5b* in the mutants *another longfin (alf)* and *pfau* (Perathoner 2014); however, the new mutation was not linked to the *kcnk5b* locus (data not shown). Given the flowing fin morphology, we named this mutant *schleier (schl)*. In addition to the dominant fin and barbel length phenotypes, the *schleier* homozygous mutant also has a novel recessive phenotype affecting stripe formation. The pattern of pigmentation along the flank of homozygotes results in broken stripes which at times resolve into spots (**Figure 1d**). Stripe pattern in the caudal and anal fins is also greatly disrupted. This phenotype is not present in *kcnk5b* mutants, which do not affect normal stripe patterning (Perathoner, et al., 2014). Thus, the *schleier* mutant is a unique locus sharing many of the same effects as *alf/kcnk5b* mutants but with a separate role in stripe patterning during late development.

**Figure 1.**
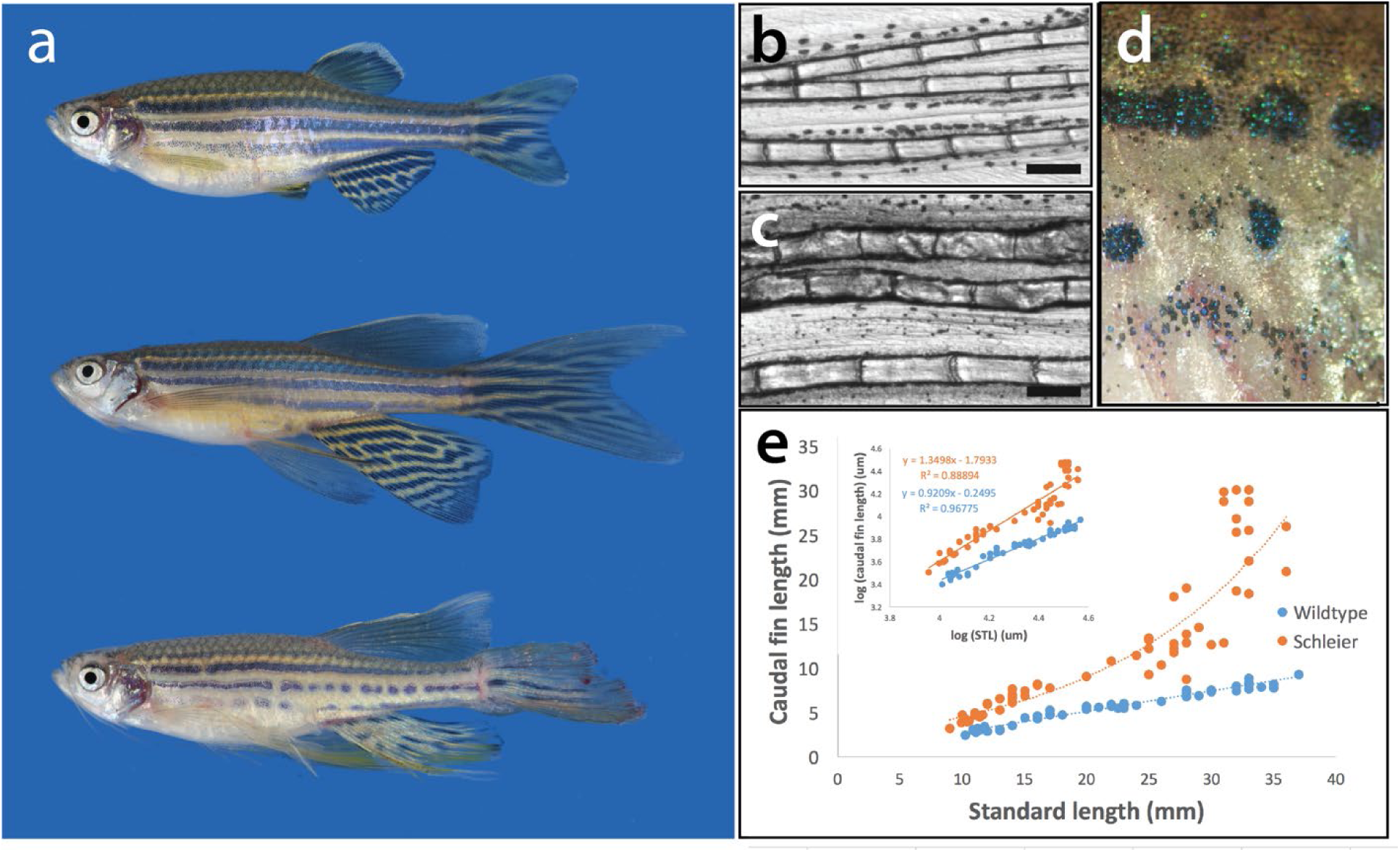
Dominant enhancer of *longfin* causes increased fin proportion through enhanced relative growth of the fin during development. **a**) Phenotype of *schleier* mutant zebrafish affects fin proportion and pigmentation of the adult zebrafish in a dose dependent manner. **b, c**) Segment length of lepidotrichia from adult caudal fins of wild-type and *schleier* heterozygous fish. Bar represents 200 µM. **d**) Close up of pigment pattern deficiencies on flank of *schleier* homozygous fish. **e**) Growth curves of caudal fin length in wild-type and *schleier* heterozygous fish as a function of standard length (STL). **f)** Log-log plot of caudal fin length vs. STL to illustrate allometric scaling of growth.

To characterize changes in fin growth, we measured caudal fin length and standard length of *schleier* heterozygotes and wildtype siblings during post-embryonic development. When compared to wild-type individuals, *schleier* heterozygotes display an increased rate of fin growth relative to standard length throughout development, leading to overall positive allometry (α=1.35) of the fins (**Figure 1e**). Such alterations in fin proportion could be caused by patterning changes in the number and/or size of segments in the dermal rays. We find that *schleier* fin rays show elongation of segment size while retaining similar numbers of segments as wild-type fish (**Supplementary Figure 2**). This finding resembles the phenotype of the *another longfin (alf) or pfau* long-finned mutants, which exhibit comparable changes in fin skeletal patterning and are caused by gain of function-alterations in Kcnk5b (Perathoner, et al., 2014). Given the phenotypic similarity between these mutants and that our previous work linked *alf*/*kcnk5b* function to bioelectric signaling in development (Daane, et al., 2018), we chose to further characterize *schleier* as it might reveal mechanistic aspects of bioelectric signaling in size regulation.

### *Schleier* mutation highlights role of vascular tone associated with altered proportionality in zebrafish fins

In addition to their overgrown fins and barbels, one notable feature of the *schleier* mutant fish is their unusual fin vasculature. Under light microscopy, caudal fins of live *schleier* fish are seen to have enlarged blood vessels and alterations in flow rate to and from the fin tip. We characterized the vascular phenotype in more depth in wild-type and heterozygous *schleier* fins in order to identify possible mechanisms underlying fin overgrowth. We crossed the *schleier* mutation into a transgenic strain, *Tg(fli1:EGFP),* in which green fluorescent protein is expressed in all vasculature (Lawson and Weinstein, 2002). Wild-type fish displayed the expected caudal fin vasculature extending along fin rays of one centrally located artery flanked by two veins in each fin ray (**Figure 2a, c**). These primary vascular elements were connected to one another by inter-vessel commissures, and to adjoining veins by inter-ray vessels. *Schleier* heterozygous fish retained overall normal vascular patterning, with significantly enlarged diameter of all blood vessels and increased number of *fli*+ endothelial cells lining vessels (**Figure 2b, d**). To investigate the functional consequence of these changes, we performed high-speed videomicroscopy on *fli:GFP* caudal fins of live wild-type and *schleier* fish. Data were analyzed to determine the velocity of individual red blood cells in veins and arteries as well as the diameter of the corresponding veins and arteries in the tissue proximal to the fin tip. The veins of *schleier* mutant fish had a consistently larger diameter relative to the adjacent artery, resulting in an increased vein:artery diameter ratio (**Figure 2e**). This structural change was accompanied by a change in the relative rates of blood flow in *schleier* vessels (**Supplementary movies 1, 2**). We quantitated this phenotype by measuring the relative vein:artery velocity, and found that relative venous flow was almost twice as slow in *schleier* caudal fins as compared to wild-type fins (**Figure 2f**). The result of this varied flow rate was apparent in gross inspection of the *schleier* mutant, as pooling of blood is often seen at distal tips of overgrown *schleier* fins and barbs (**Figure 2h)** but not in wild-type fins (**Figure 2g**).

**Figure 2.**
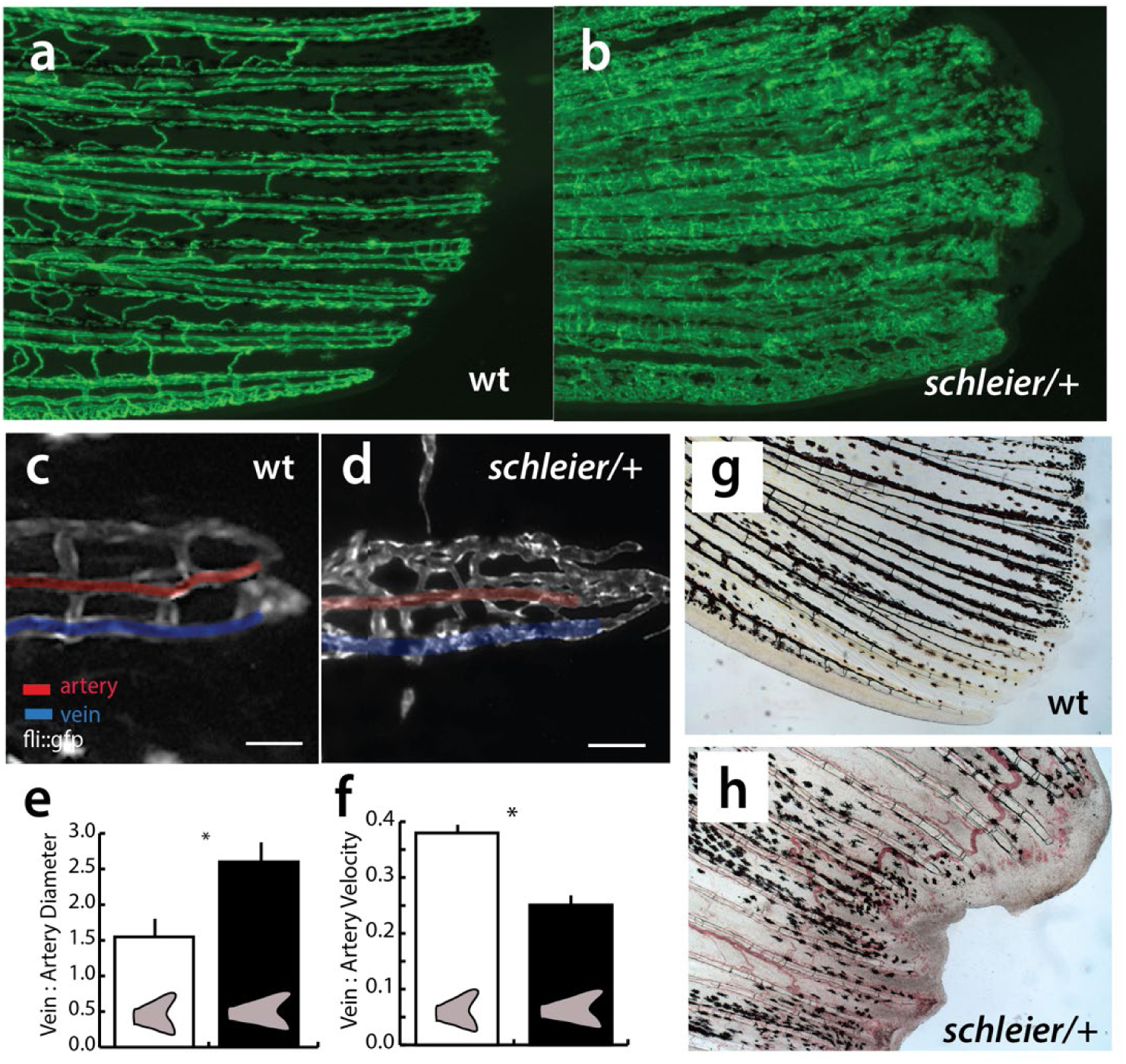
*Schleier* mutation highlights role of vascular tone associated with altered proportionality in zebrafish fins. **a, b**) Pattern of vascularization in caudal fins of adult wild-type (wt) and *schleier* heterozygous fish visualized by *Tg(fli:gfp)* transgenic expression of endothelial cells. **c, d**) Close up of caudal fin tip showing venous and arterial pattern variation in the *schleier* mutant associated with increased number of endothelial cells. Veins and arteries pseudo-colored blue and red, respectively; scale bar equals 100 µM. **e**) Measure of vascular dilation in *schleier* caudal fin tips. Mean +/- standard error of mean (SEM) is shown. *, p<0.05. **f**) Quantitation of vascular flow in fin tip in arteries compared with veins. *, p<0.05. **g, h**) Excessive vascularization and blood pooling in the *schleier* mutant fin tip (h) compared with wild-type fin tip (g).

### *schleier* is caused by mutation of *slc12a7a* encoding potassium chloride cotransporter *kcc4a*

As the *schleier* mutation originated from a screen on homozygous *lof(dt2)* background, the *schleier* locus will retain a *lof*-specific haplotype in the region of the mutation. Outcross of the original F1 mutant revealed that the mutation was linked to the *lof* locus but could be recombined away from it (39/134 outcross progeny). We then used homozygosity mapping and massively parallel sequencing (methods previously defined in (Bowen, et al., 2012)) to link the *schleier* mutation to the *slc12a7a/kcc4a* gene on chromosome 2. To screen for potential causative mutations, we used whole-genome-sequencing-generated databases of variant SNPs for the *lof* background (Bowen, et al., 2012) to identify unique changes that were present in *schleier* and never seen in *lof*. From this analysis, we identified two SNPs located in *slc12a7a/kcc4*a on Chromosome 2 that were unique to the *schleier* mutant sequence. One SNP was located in the 3’ UTR of a putative alternative transcript, while the other SNP led to a missense mutation (C583Y) of a conserved residue in exon 14 in the primary transcript of the gene. Through outcrossing the mutant, we were able to recover recombinant fish that had fin overgrowth and retained only the C583Y mutation.

*Slc12a7a* is predicted to encode a potassium chloride co-transporter Kcc4a. Given our previous finding that *kcnk5b*, a potassium channel, is involved in regulating proportion and has an overgrowth phenotype similar to *schleier*, *slc12a7a* was a strong candidate. We identified a single unique missense mutation in the *slc12a7a* gene, C583Y, present in all *schleier* mutants and absent in wild-type siblings (0/56) (**Figure 3a**). Kcc4a protein that contains the C583Y mutation is likely not functional, as the mutation eliminates a cysteine residue predicted to form a critical disulfide bond (DiANNA analysis; (Ferre and Clote, 2005)).

**Figure 3.**
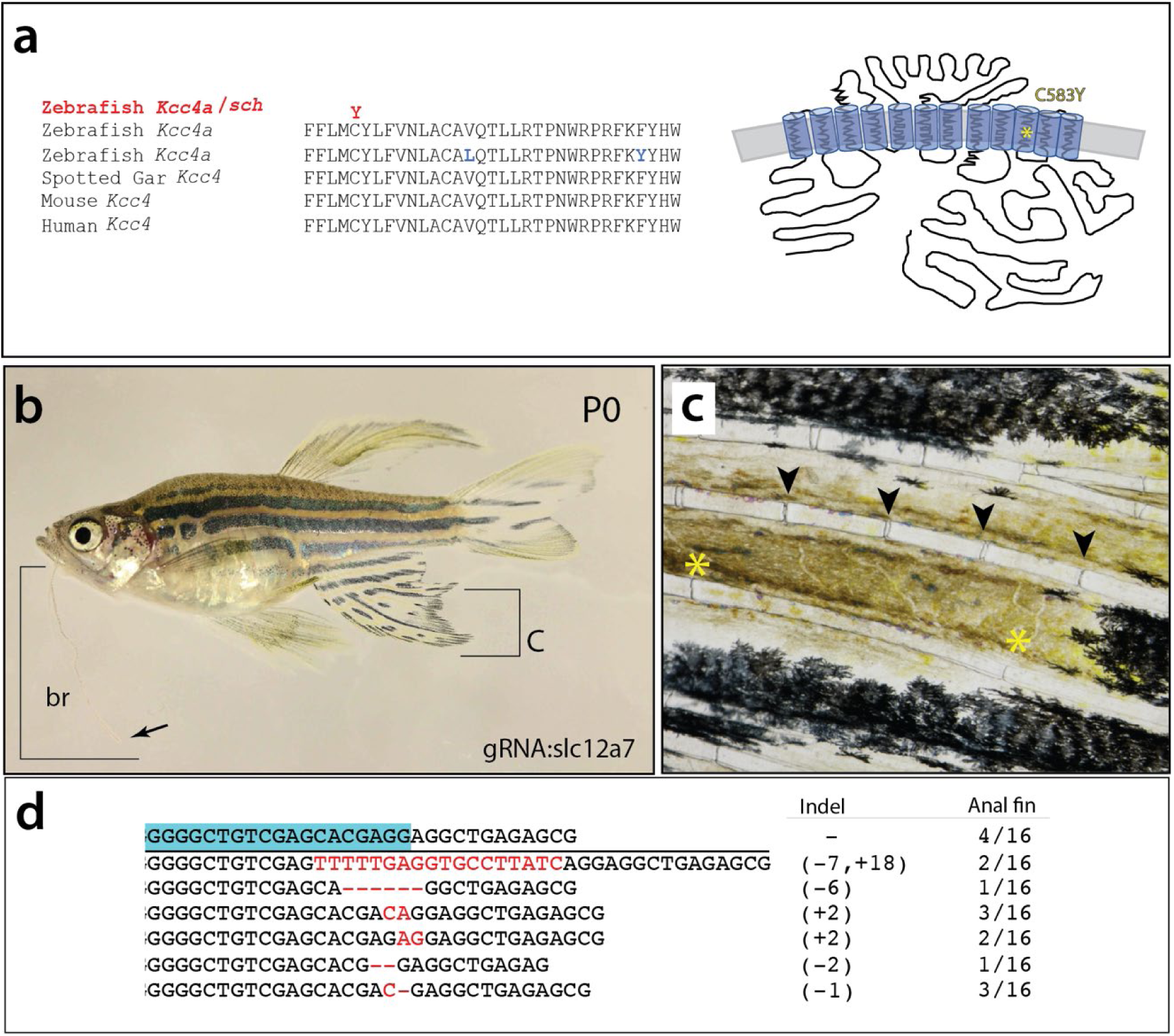
*Schleier* is caused by mutation of *slc12a7a* encoding potassium chloride cotransporter Kcc4a. **a)** Multispecies alignment of predicted coding sequence of *slc12a7a* showing position of altered residue in Kcc4a in the *schleier* mutant. Schematic representation redrawn after (Marcoux, et al., 2017). **b**) Adult wildtype P0 zebrafish with somatic clones harboring deletions of *slc12a7a/kcc4a*. Arrow points to top of barbel; bracket shows extent of growth. **c**) Close up of anal fin labeled in (b) showing elongated segments of the rays (joints marked *); normal branch pattern observed just above (arrowheads). **d**) Sequence of deletions identified in affected region of fin and frequency (12/16); blue box, sequence of the guide RNA used in targeting.

To test whether mutations in *slc12a7a/kcc4a* could result in fin overgrowth, we designed specific guide RNAs (gRNAs) targeted against the third exon of the gene that should affect all predicted isoforms of Kcc4a. These guides were injected with Cas9 mRNA into wild-type embryos. Strikingly, adults from injected embryos (P0) showed broad proportional overgrowth of fins and barbels (**Figure 3b**). Additionally, similar to homozygous *schleier* mutants, an alteration of pigmentation stripe pattern was seen in the anal and caudal fins (**Figure 3b**). The observed phenotypes were localized to specific areas of the fin and likely due to mosaicism in injected animals. Overgrown areas of fins revealed elongation of fin ray segments similar to that seen in *schleier* (**Figure 3c**). Genotypic analysis of these areas of overgrowth fin showed deletions in exon 3 of *slc12a7a/kcc4a* at high percentage (12/16; 75%; **Figure 3d**). The majority of the mutations detected (11/12; 92%) were predicted to cause a frameshift in the translated protein resulting in a premature stop codon shortly thereafter.

Zebrafish have two *slc12a7/kcc4* paralogues that share 74% amino acid identity. By *in situ* hybridization, *slc12a7a/kcc4a* transcript was detected at modest levels in wildtype intra-ray and inter-ray fin mesenchyme and epidermis (**Figure 4a**). *Schleier* heterozygous fin tissue had decreased transcript expression in both fin epidermis and fin mesenchyme (**Figure 4b**). Using antibodies against Kcc4 paralogues, we detected a single band of expected size when used on a Western blot of zebrafish tissue extracts (data not shown). As the target antigen is present in both Kcc4a and Kcc4b proteins, the antibody is predicted to recognize both paralogues in the zebrafish. The expression pattern of the Kcc4a/b proteins in fin cryosections showed Kcc4a/b protein expression in a pattern consistent with *in situ* hybridization specific for *slc12a7a/kcc4a*. Kcc4a/b were also detected in the epidermis with heightened punctate expression in specific cells or regions. In *schleier* fin mesenchyme and epidermis, immunofluorescence revealed decreased amount of Kcc4a/b protein as compared to wild-type fin (**Figure 4c and 4d**, arrowheads).

**Figure 4.**
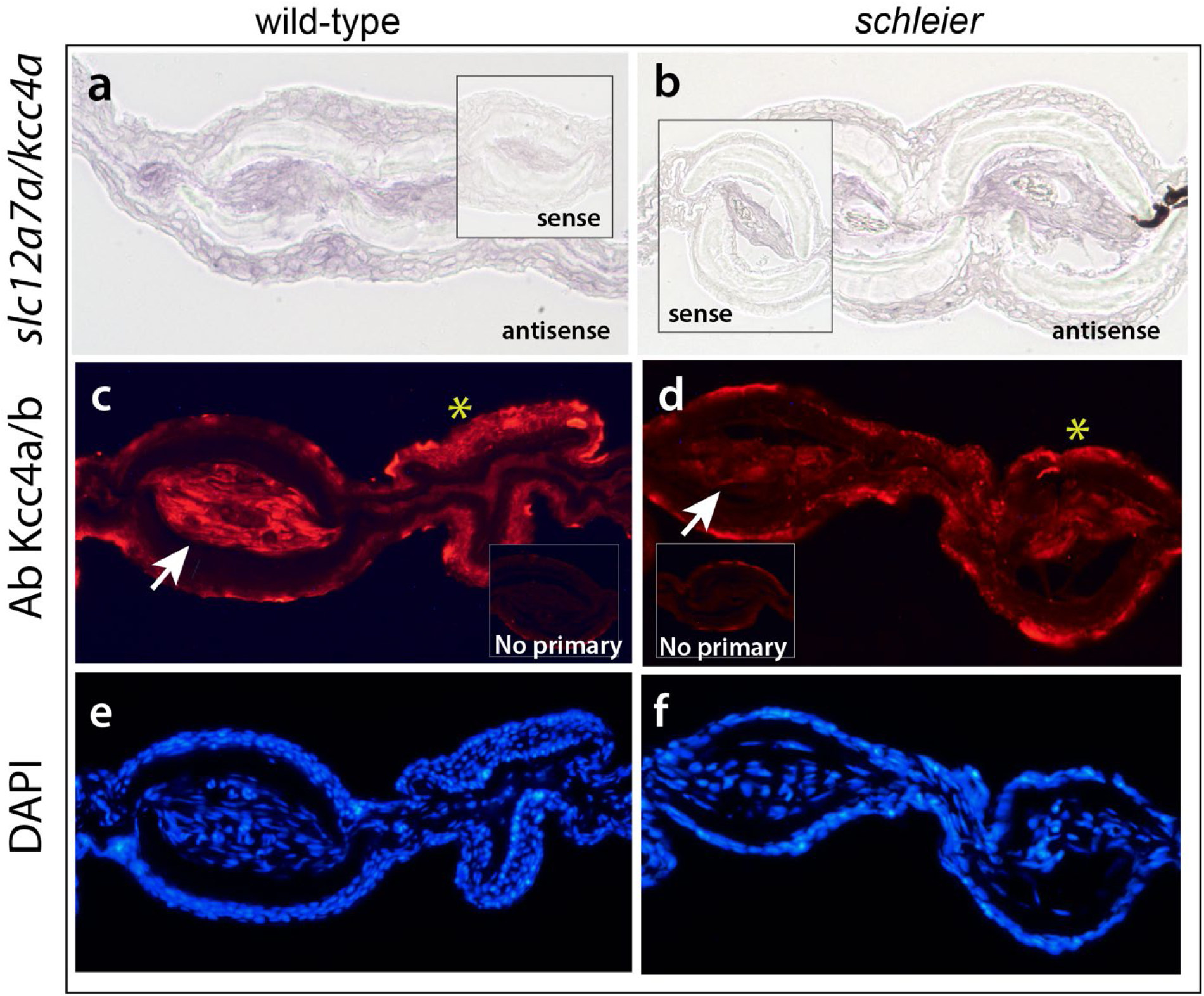
The *schleier* C583Y mutation causes decreased Kcc4a expression in fin mesenchyme. **a, b)** Transcriptional regulation *of slc12a7a/kcc4a* in adult caudal fins of wild-type (a) and *schleier* mutants (b). *Inset*, signal from control *in situ* probes on serial sections. **c-d)** Immunofluorescence of antibodies against Kcc4 in adult caudal fins. Arrow points to mesenchyme of ray, asterisk, epidermis of fin. *Inset,* signal from control sections hybridized with secondary antibody only. **e, f**) DAPI counterstaining of fin sections.

### *Schleier* C583Y mutation causes transporter loss-of-function

Our previous work on the *kcnk5b/alf* long-finned mutant established that the *alf* mutation caused increased potassium transport. As the Kcc4a protein also mediates potassium transport, and the *schleier* mutant has strong phenotypic similarity to the *alf* mutant, our *a priori* expectation was that the *schleier* mutation would also cause increased protein activity. However, our gene targeting of *slc12a7a/kcc4a* indicated that loss-of-function mutations in the gene were sufficient to confer a long-finned phenotype. Given this unexpected result, we decided to determine the effect of the *schleier* C583Y mutation on the transporter activity of the Kcc4a protein.

cRNAs encoding wild-type or Kcc4a C583Y mutant protein were injected singly or as a mixture into *Xenopus laevis* oocytes under hypotonic conditions. Notably, the presence of the C583Y mutation resulted in a 90% decrease in transport activity (WT, 11686.7 ± 1555.3 vs C583Y, 968.1 ± 392.7 pmol/oocyte/h, p<0.001; **Figure 5a**), without modifying the total protein expression (**Figure 5**, **inset**). When co-expressed with C583Y, wild-type cotransporter function was significantly affected at the higher protein concentration tested (WT, 10110.1 ± 1256.3 vs WT:C583Y (0.2 µg/µl), 6785.5 ± 729.3 pmol/oocyte/h, p<0.05) (**Figure 5b**). Based on these observations, we conclude that the C583Y mutation causes loss of ion transport function in the KCC4a protein.

**Figure 5.**
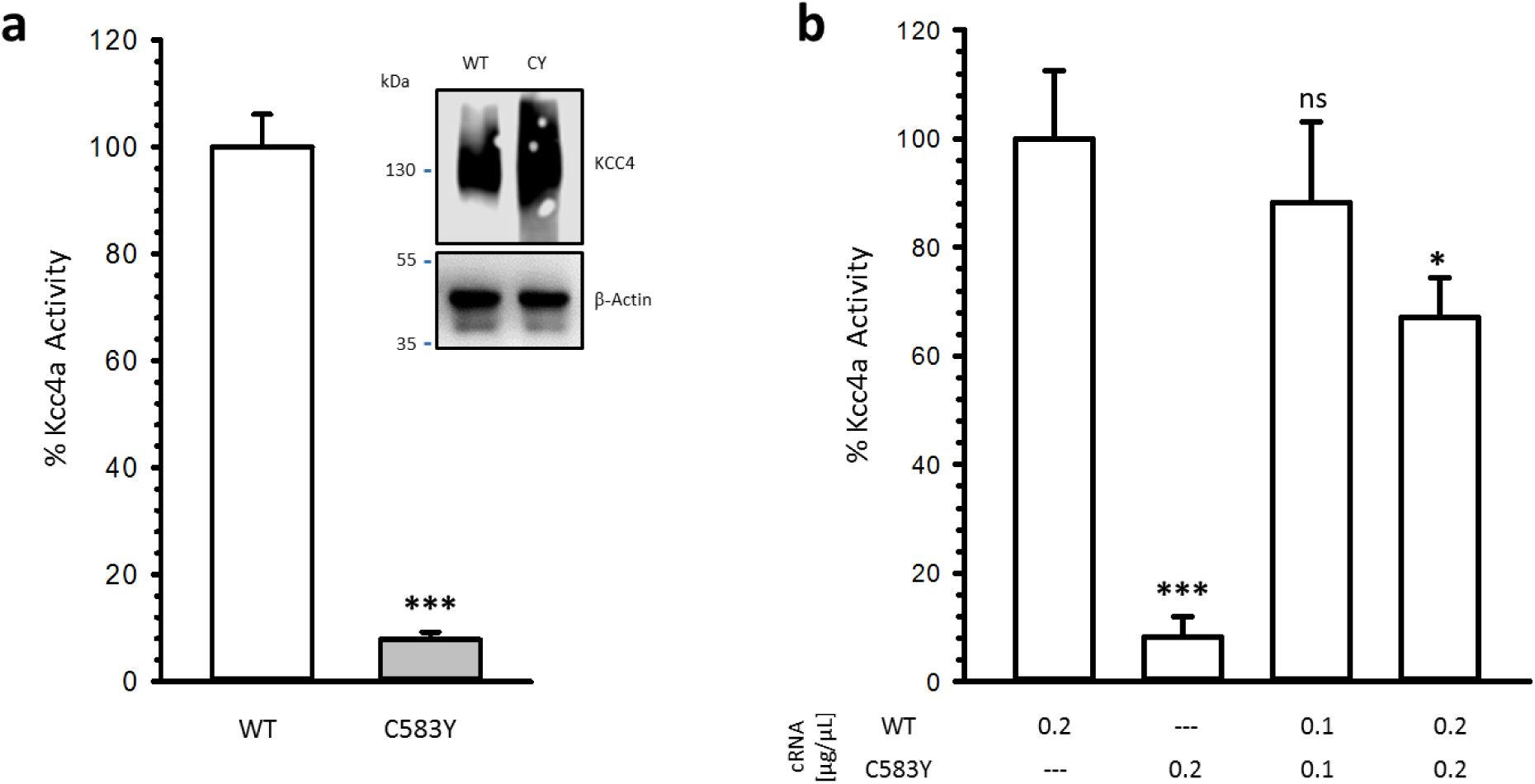
The effect of the C583Y mutation results in disruption of co-transport function of Kcc4a. **a)** Percentage of activity was determined as the Cl^−^-dependent, Rb^+^ cotransport by Slc2a7a/Kcc4a tested in *Xenopus oocytes* through expression of wild-type (WT) and C583Y Kcc4a under hypotonic conditions. *Inset* shows protein expression from total extracts. **b)** Percentage of Kcc4 activity co-expressed with different concentrations of C583Y mutant as shown. For both **a** and **b**, mean +/- SEM are shown. n.s., not significant; * p<0.05; * p< 0.0001 calculated using a student’s T test in comparison to wildtype Kcc4a. Results presented are the summed activity of three separate experiments; n=30 oocytes per group.

### In-frame deletions in the *slc12a7a/kcc4a* gene cause alterations in fin size

The effect of Crispr-Cas induced mutations in s*lc12a7a/kcc4a* in wild-type fish showed that mosaic inactivation of the gene was sufficient to cause fin overgrowth. However, it remained unknown which particular mutant alleles were capable of causing this phenotype. To address this question, we crossed mosaic P0 fish to wild-type fish to establish individual lines of F1 progeny heterozygous for unique alleles of *slc12a7a/kcc4a*.

The F1 progeny displayed a range of phenotypes, from phenotypically wild-type to fish with long fins and extended barbels (**Figure 6c**). Genotyping of phenotypically wild-type F1 fish revealed that some of these fish were heterozygous for deletions predicted to cause premature stop codons in exon 3 (**Figure 6a**). Upon translation, such mutations would produce a truncated Kcc4a protein that was 99 amino acids in length, as compared to the 1117 amino acid wild-type protein; such a protein is probably nonfunctional and/or unstable. Based on the normal fin and barbel phenotypes observed in fish heterozygous for truncation alleles, we concluded that 50% reduction in Kcc4a protein was not sufficient to cause fin and barbel overgrowth.

**Figure 6.**
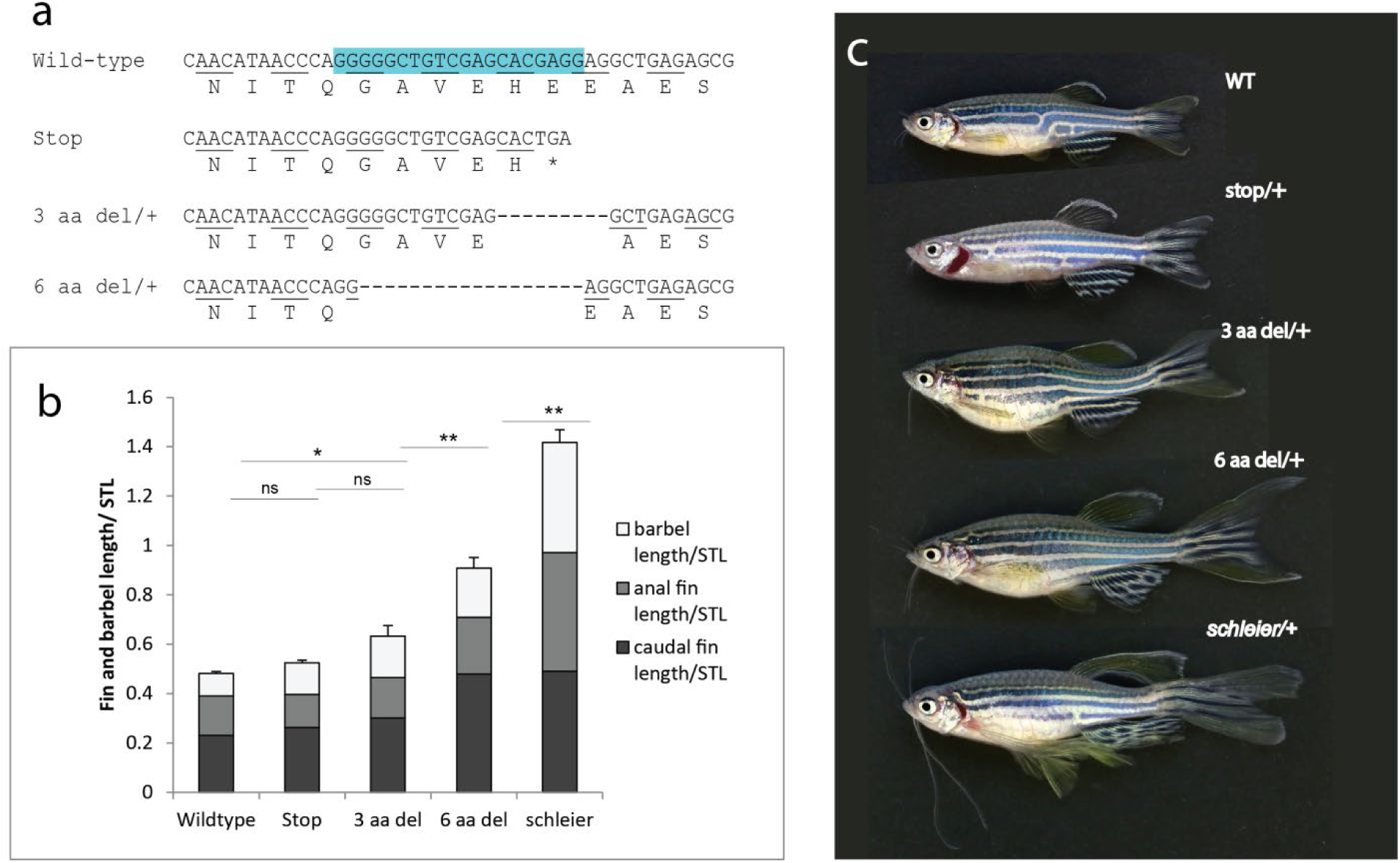
Allelic titration of *slc12a7a*/*kcc4a* expressivity revealing necessity and sensitivity to levels of Kcc4a during development. **a)** Identified deletions in F1 progeny and predicted protein alteration; blue box, sequence targeted by guide RNA. **b)** Tabulation of relative length of fins and barbels in F1 progeny heterozygous for each genotype; mean +/- SEM is shown. n.s., not significant;* p< 0.05; ** p<0.01 calculated using a one-way ANOVA followed by a Tukey’s multiple comparisons test; n=6-19 fish per genotype. **c**) Representative pictures of individuals heterozygous for different *slc12a7a/kcc4a* alleles.

We also genotyped long-finned F1 fish to identify mutations that caused fin overgrowth and found that these fish were heterozygous for small in-frame deletions or insertions in exon 3 (**Figure 6a**). Two such alleles were isolated from multiple independent founders. One allele contained a three amino acid deletion and caused only mild phenotypic changes such as long barbels (**Figure 6c**). A second allele caused a six amino acid deletion that resulted in significantly elongated barbels and long caudal and anal fins; this mutant also exhibited disrupted stripe pattern in the anal fin (**Figure 6c**). These phenotypes were similar to those seen in the *schleier* mutant but with decreased severity. Interestingly, the size of the deletion correlated with expressivity of the phenotype (**Figure 6**) suggesting that Kcc4a is quite sensitive to dose of expressed, but nonfunctional variants.

### Epistasis of potassium channel signaling mediating reveals common pathway of growth regulation

We have previously demonstrated that activating mutations in the potassium channel *kcnk5b* caused patterned overgrowth of fins and barbels, and that absence of *kcnk5b* had no effect on fin phenotype(Perathoner, et al., 2014),(Daane, et al., 2018). Given the similarity between the *kcc4a* and *kcnk5b* overgrowth phenotypes, we hypothesized that they might work in a similar pathway or act in parallel to regulate growth.

We decided to test for interaction between the *slc12a7a/kcc4a* and *kcnk5b* genes. We designed two guide RNAs to target exon 1 of *kcnk5b* and co-injected them into *schleier* heterozygous embryos and their wild-type siblings. Injected P0 fish were grown to maturity, genotyped, and assessed for their fin and barbel phenotypes. Clonal analysis on fin tissue verified that the guide RNAs successfully introduced deletions and/or insertions at the targeted sites (**Figure 7d**). As would be expected based on our earlier work (Perathoner, et al., 2014),(Daane, et al., 2018), targeted inactivation of *kcnk5b* had no effect in a wild-type genetic background (**Figure 7a**). However, CRISPR-mediated inactivation of *kcnk5b* in a heterozygous *slc12a7a/schleier* background produced fish with reduced fin and barbel length (**Figure 7c**). Barbel length was nearly restored to wild-type, while anal fin length was significantly shorter than in *slc12a7a* heterozygotes (**Figure 7e, 7g**). The average length of individual anal fin segments was also shorter in *slc12a7a +/-*; *gkcnk5b* fish, and the shape of the caudal fin was narrower and less billowy than the veil-like *schleier* mutant phenotype (**Figure 7f, 7c**). Unlike the regional overgrowth caused by upregulation of *kcnk5b*, the phenotype of *slc12a7a/kcc4a +/-*; *gkcnk5b* fish appeared to be systemic in nature, affecting whole structures and exhibiting consistent phenotypes in all fish.

**Figure 7.**
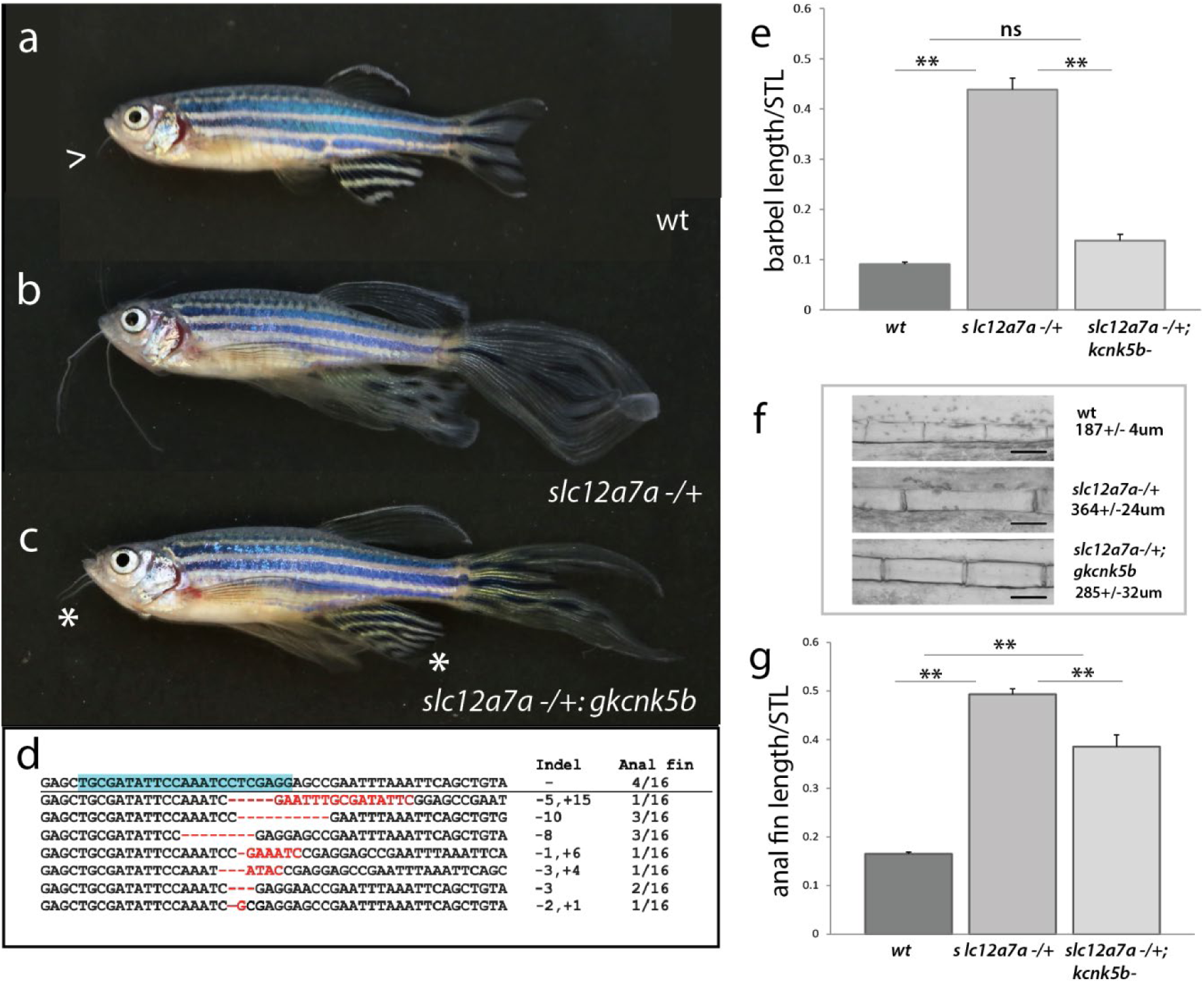
Regulation of size in *schleier* requires functional *kcnk5b.* **a-c**) Representative pictures of growth phenotypes of zebrafish with altered potassium channel function. **a)** Wild-type adult zebrafish; arrowhead, end of barbel. **b**) *schleier* heterozygote. **c**) *schleier* heterozygote fish harboring somatic deletion clones within *kcnk5b*, (*gkcnk5b*); asterisk demarcates reduction of overgrowth phenotype of the barbel and anal fins. **d**) Recovered deletions observed in *kcnk5b* from reverted anal fins in affected mutants. Blue highlight = targeted sequence. **e-g**) Reversion of *schleier* overgrowth phenotypes after abrogation of *kcnk5b* function. **e**) Relative barbel length. **f**) Segment size of fin rays. **g**) Relative anal fin length. For **e-g**, mean +/- SEM is shown for each panel. n.s., not significant;** p<0.01 calculated using a one-way ANOVA with a Tukey’s multiple comparisons test; n=7-24 fish per genotype.

## Discussion

Research into bioelectric signaling has seen a recent boom, revealing essential roles of electrochemical fields and ion channel activity during development and regeneration (McLaughlin and Levin, 2018). Importantly, many of these functions can be attributed to non-excitable tissues with changes reflected in consistent shifts of resting membrane potential of cells and across tissues. Genetic models of ion channel mutations often have very specific neurological or physiological phenotypes. Of the channels discussed in this work, both Kcnk5b (TASK2) and Kcc4 knockout mice have kidney defects due to alteration in ionic regulation in the thick ascending arm of the kidney (Warth, et al., 2004; Barriere, et al., 2003). Kcc4 KO mice additionally have hearing defects (Boettger, et al., 2002). However, neither of the mouse knockouts provides insight into the potential role of ion channels in growth and patterning, and it remains unclear whether bioelectric signaling plays a consistent role in these processes. Clinical presentations of ion channelopathies retain this bias towards phenotypes caused by changes in excitable cell populations such as muscle, nerve or endocrine tissue (Felix, 2000).

The Kcnj2/Kir2.1 protein provides one of the rare examples in which changes in channel function lead to patterning defects and dysmorphologies (Dahal, et al., 2017; Dahal, et al., 2012). Analysis of Kcnj2 in *Xenopus* has shown that modification of resting membrane potential across early ectoderm can alter craniofacial development, consistent with the spectrum of clinical phenotypes caused by mutations in this channel (Adams, et al., 2016; Donaldson, et al., 2004). Zebrafish mutants provide other prominent examples that begin to illuminate the extent of potassium channel regulation in growth and development, such as Kcnj13 in patterning pigment stripes (Iwashita, et al., 2006) and Kcnk5b in regulating fin size regulation (Perathoner, et al., 2014). These mutants are for the most part gain-of-function and thus can be considered supra-physiological cases not resembling ‘normal’ mechanisms of development. Here, we describe a functional role for *slc12a7a/kcc4a* in regulating size and growth of the zebrafish fin through diminishing the fraction of functional channels (**Supp. Fig. 2**). These data support the essential nature of bioelectric signaling in the regulation and development of proportion.

### Genetic basis of Kcc4a function in size regulation

Kcc4 has two paralogues in zebrafish, *kcc4a* and *kcc4b*. Our expression data suggest that the two paralogues and their encoded isoforms have overlapping patterns of expression. *slc12a7a/kcc4a* appears to be expressed primarily in fin mesenchyme and epidermis. Its paralogue, *slc12a7b/kcc4b*, may also have an epidermal expression pattern, but further analysis would be required to support this hypothesis. For both ISH and immunofluorescence detection methods, presence of the *schleier* C583Y mutation resulted in decreased expression of transcript and protein in the fin mesenchyme and epidermis.

Our data identify transport activity by Kcc4a as essential for normal fin size regulation. We observe that the *schleier* mutation in *slc12a7a* causes loss of transport function. However, fish heterozygous for a premature stop mutation in *slc12a7a/kcc4a* have normal fin size (**Figure 6**); thus, the mutation is not haploinsufficient. We propose that the *schleier* mutation has a dominant negative effect on the Kcc4a protein. Given that the protein functions as a membrane bound dimer (Bergeron, et al., 2011), mutations such as *schleier* that result in the formation of full-length, though inactive, protein should cause a greater decrease in the levels of functional Kcc4a dimer than loss of one allele (**Supplementary figure 3**). Consistent with this, we observed that heterozygosity for in-frame deletions and the *schleier* C583Y mutation, but not heterozygosity for a null mutation, resulted in fin and barbel overgrowth. Interestingly, the phenotypic effects of Kcc4a gain-of-function mutations are scaled to the size of the in-frame deletions and can be titrated (**Figure 6**).

Identification of the role of *kcc4a* in size regulation may provide a useful *in vivo* model for investigation of the molecular properties of the Kcc4a protein. Through genome editing, functional requirements for specific domains of the protein and putative regulatory sites can assessed *in vivo*. To add further complexity, the Kcc4 protein has been shown to form heterodimers with other KCC and NKCC family members via their C-terminal domains (Simard, et al., 2007). Overlapping expression of these heterodimers *in vivo* might buffer against fluctuations in potassium transport caused by loss of a single *slc12a7a*/*kcc4a* allele. It would be of interest to investigate whether inactivation of the paralogue *slc12a7b/kcc4b* or of other *kcc/nkcc* family members increases the severity of the observed phenotype.

### Dose dependent functions and threshold responses

The *schleier* mutation in *slc12a7a/kcc4a* causes a dominant increase in the relative proportion of the fins and barbs as well as a recessive pigment phenotype affecting patterning of the adult stripes. Among zebrafish with mutations affecting conductivity, *schleier* is the first mutant to share both fin and pigment phenotypes. These pleiotropic effects suggest the mutated Kcc4a channel may oligomerize with other KCC or NKCC channels, including Kcc4b, to inhibit or modify their function (Simard, et al., 2007). Interestingly, the *schleier* pigment phenotype closely resembles the phenotype of fish with mutations in *connexin 41.8* (Irion, et al., 2014; Watanabe, et al., 2006) but not those with mutations in *kcnj13* (Iwashita, et al., 2006). Thus, Kcc4a may be tied to the regulation of bioelectric signaling through Cnx41.8 mediated cell connectivity in pigment patterning and through Cnx43 in development of fin length (Perathoner, et al., 2014; Iovine, et al., 2005). The presence of the pigment phenotype in homozygous *kcc4a/schleier* mutants but not heterozygotes could reflect differential sensitivity of growth versus pigment pathways to changes in *Vmem*, or alternatively, could reflect a dominant negative effect of the *schleier* mutation on other signaling processes and other channels.

How bioelectric signals are transduced into a growth response in the fin, and their specificity, remains an open area of investigation. Changes in membrane polarization may alter influx and balance of critical intracellular signaling ions like calcium, resulting in changes to the transcriptional program and cell proliferation. Alternatively, or in addition to this, changes to the overall ion concentration in a tissue environment may directly alter the properties of local cells and tissues. Potassium is known to be a critical regulator of vascular tone. Inactivation of *kcc4a* in fin mesenchyme might result in increased potassium in extracellular fluid adjacent to the fin blood vessels, resulting in vasoconstriction of the arteries. In support of this, we observed that fins with inactive *slc12a7a*/*kcc4a* had increased blood flow to the fin tip. This blood pooling at the distal margin of the fin is accompanied by persistence of a capillary nexus and is present in both *slc12a7a*/*kcc4a* and *kcnk5b* long-finned mutants. The increased arterial blood flow was accompanied by slower return transit through fin veins, which were grossly enlarged to accommodate the larger volume (**Supplementary movie 2**). These anatomical changes could prolong exposure to circulating growth factors; thus, ion-mediated changes in vascular tone might provide one means via which these mutations affect growth.

### Genetic epistasis among potassium channels and regulation of coordinated growth and size

Our data suggest that potassium homeostasis within cells and across tissues may be a key regulator of size in the zebrafish fin. In previous work, we demonstrated that gain-of-function mutations in the potassium channel protein Kcnk5b result in fin and barbel overgrowth. Here, we show that mutations in Kcc4a, another protein involved in potassium transport, produce a similar long-finned phenotype. The phenotypes of *schl/kcc4a* and *alf/kcnk5b* closely resemble each other, suggesting they may regulate similar pathways. The expression pattern of Kcc4a in the fin mesenchyme is similar to the tissue we have shown is responsible for *kcnk5b* growth regulation in transplants (Perathoner, et al., 2014); thus, these factors are likely operating in the same cells. Our findings further suggest that the Kcc4a and Kcnk5b proteins work in concert, and that alterations in the function of either component can increase growth. The *kcnk5b* long-finned mutations cause increased potassium transport, while the *schleier* mutation causes a loss-of-transport function. Although deletion of *kcnk5b* fails to have a growth phenotype, we have previously shown that the function of this channel was necessary for enhanced growth caused by FK506-mediated inhibition of calcineurin (Daane, et al., 2018). Here, we show that *kcnk5b* is also required for the effect of the *schleier* mutation in Kcc4a. Whether this genetic relationship is additive or epistatic remains an open question. We find that unlike overexpression of *kcnk5b* or loss of *slc12a7a*/*kcc4,* which act in a local manner in the fin causing outgrowth, loss of Kcnk5b seems to affect size regulation of the fins overall, suggesting a potential systemic role in suppression of growth. As both channels are expressed in the kidney and play roles in regulating hypertension, this may be a facet of regulation for further investigation.

### Core components of growth regulation mediated by integration of potassium channel function

The alteration in the net flow of ions within a tissue will have a direct effect on the resting membrane potential of a cell, *Vmem*. Consistent with this fact, we have shown that gain-of-function mutations in Kcnk5b lead to hyperpolarization of *Xenopus* oocytes (Perathoner, et al., 2014). We favor an additive hypothesis for regulation of growth signals by potassium channel regulation, with the effects of each gene summated in the resting membrane potential of a cell (**Figure 8a**). In this model, channels such as Kcc4a and Kcnk5b integrate environmental stimuli as well as systemic regulators of growth into classic signaling pathways in tissues, coordinating diverse signals into a bioelectric response (Daane, et al., 2018; McClenaghan, et al., 2016; Bagriantsev, et al., 2011). We propose that multiple inputs affect potassium ion concentration and contribute to a common bioelectrical signaling response in the zebrafish fin. This system is sensitive even to slight alterations in protein function, as seen in the *slc12a7a/kcc4a* CRISPR alleles. Classical growth factors may also provide input to this network, such as Igf1, an important paracrine regulator of relative proportion that acts in part to control coordinated growth within tissues (Yakar, et al., 2018). Notably, Igf1 has been found to upregulate Kcc3 and Kcc4 activity in cancer cells, suggesting it can mediate activity of this proposed network (Hsu, et al., 2007; Shen, et al., 2001). The potassium channel *kcnk5b* may play a core role in modulating the output of variations in *Vmem*, as it is needed for overgrowth in multiple contexts (Kcc4 inactivation, calcineurin inhibition (Daane, et al., 2018)) and can itself lead to overgrowth when locally overexpressed or activated (Perathoner, et al., 2014).

**Figure 8.**
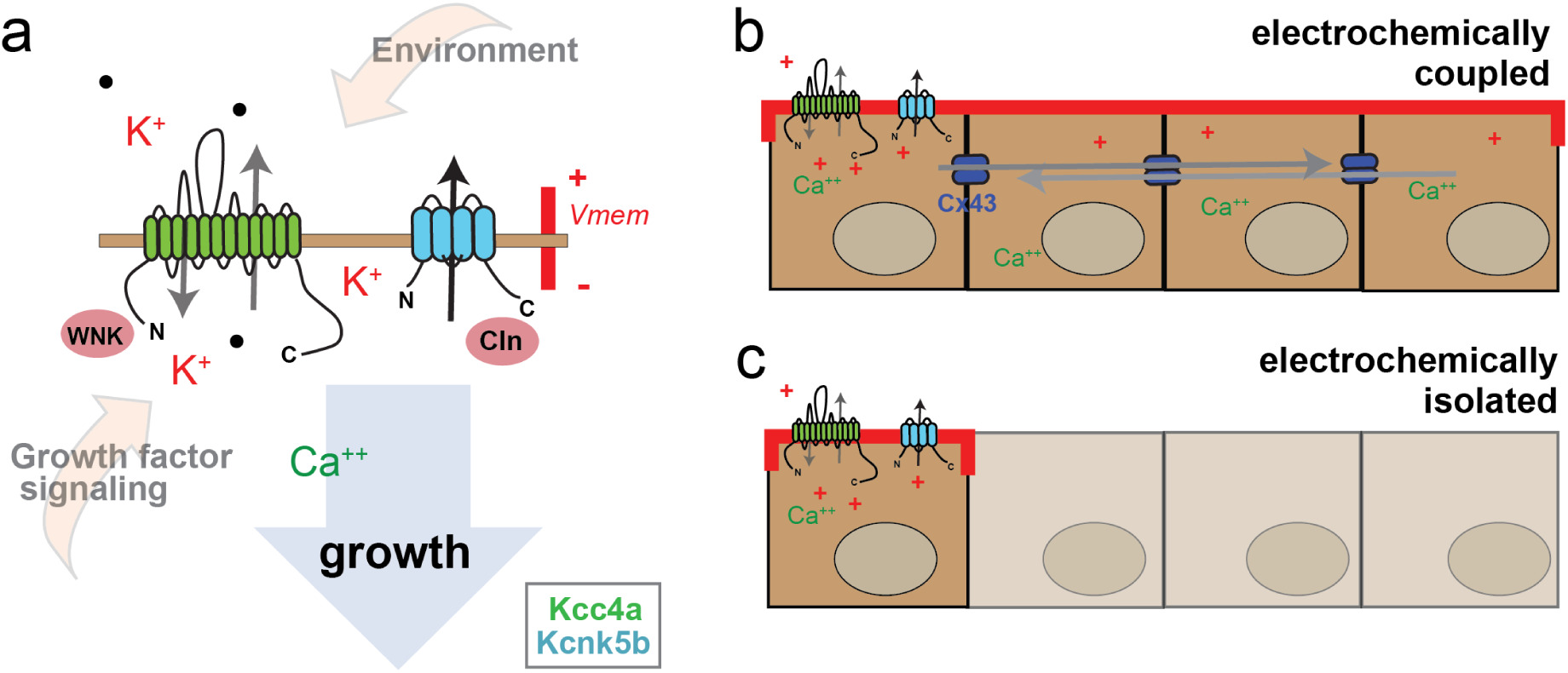
Model of bioelectrical signal integration during growth of the zebrafish fin. Multifaceted integration of environmental and classical secondary signal messengers in regulating resting membrane voltage by potassium channel activity. **a**) Regulation of ionic conductance across a plasma membrane through additive inputs of Kcc4a and Kcnk5b channel regulation. Kcc4 is electroneutral, transporting K+ and Cl− (•) ions across membrane; at normal resting potential, K+ flux is normally outwards. The action of KCC channels can be regulated by cytoplasmic factors such as WNK kinases allowing integration with other signaling pathways. Kcnk5b is an outward leak channel gated in part through environmental conditions such as pH, as well as internal signaling factors binding to the C-terminus. Calcineurin (Cln) is a key signaling factor that links Kcnk5b function to canonical signaling pathways (Daane, et al., 2018). We suggest that the combined output of Kcc4a and Kcnk5b alters resting membrane potential, and that this ionic modulation may serve as a broader readout of environmental changes and of levels of intracellular secondary messengers resulting from growth factor signaling. As Kcnk5b is necessary for expression of growth effects of Kcc4a and calcineurin inhibition, we posit that it acts as translator of changes in membrane potential to calcineurin-mediated growth. **b-c**) Model of electrocoupling of cells and potential for coordinated growth in development. **b**) A shift in membrane resting potential can have effects that extend across cells and thus coordinate potentially noisy developmental signals. This regulation is mediated in the developing and regenerating fin by gap junctions comprised of Connexin-43. This electrocoupling leads to cell non-autonomous effects in growth as seen in loss of *slc12a7a/kcc4a* or *kcnk5b* expression. **c**) Lack of cell electrocoupling; signals would be predicted to be restricted and less coordinated.

In this bioelectric signaling model, we would predict that membrane polarity is interpreted through steady state alterations in net *Vmem* within tissues (**Figure 8**). Each potassium channel type will have specialized functions and will modify *Vmem* with different sensitivities to changes in local ionic potential as well as secondary signaling capabilities. Some channels may provide different phenotypic effects depending on their biological context. For example, our data demonstrate that loss of Kcc4 activity results in increased growth of the fin; however, clinical data indicate that increased Kcc4 activity correlates with invasiveness in human cancer cells (Brown, et al., 2018; Zhou, et al., 2018; Chen, et al., 2009). These seemingly conflicting observations lend support to the concept that alterations in resting *Vmem* are key to regulating growth and coordinated cell interactions in tissues during development and homeostasis (Levin, 2014). In the context of the zebrafish fin, our model predicts that decreasing the potassium gradient across mesenchymal and epidermal cells leads to hyperpolarization of resting membrane potential and promotes growth (**Figure 8a**). Manifestation of this growth response within cells of the developing fin also requires intracellular communication through gap junctions (**Figure 8b, c**) (Perathoner, et al., 2014). An extension of this model is that reduction or elimination of this connectivity may underlie the loss of coordinated growth in neoplastic tissues (Litan and Langhans, 2015; Lobikin, et al., 2012).

## Conclusions

Here we find that potassium channel function and bioelectric signaling are necessary for the normal establishment of proportion and size in development. We posit a model in which growth of the fin is modulated by relative changes in *Vmem*. This signal is translated into growth in part through the action of Kcnk5b, which may act as a central rheostat to modulate calcium-dependent growth (Daane, et al., 2018). Application of this bioelectric model to understand neoplastic growth and migration is pertinent given that many tumor types exhibit increased expression of potassium channels from diverse protein families and with different conductive and gating properties (Lang and Stournaras, 2014; Prevarskaya, et al., 2010; Wang, 2004). Further analysis of regulators of this genetic pathway may reveal the checks and balances of tissue formation, homeostasis, and how coordination of growth is regulated during development and regeneration.

## Acknowledgements

The authors would like to thank prior work by Simon Perathoner, Iris Koch, and Ines Gehring on the *schleier* mutant and its isolation. JL thanks Amanda Bettle and Beth Amaral for fish husbandry. This work was partially supported by funding by NIH R01HD084985 to MPH. KK is supported by NIH 1 R01 NS109358-0, Simons Foundation, March of Dimes, and the Hydrocephalus Association. The funding agencies had no input on the study design nor presentation of the data within.

## Author Contributions

JL, KK, and MPH designed experimental strategy in this work; JL, KK, JD, DP, LE, MB, AM, GG performed experiments supporting the data presented. JL and MPH wrote first and JL, KK, and MPH were involved with subsequent drafts and revisions of the paper.

## Competing Interests

The authors have no competing interests regarding the data presented in this work.

## Methods

### Husbandry

All fish strains used were housed and maintained as described in (Nusslein-Volhard and Dahm, 2002) and performed in accordance with IACUC guidelines at Boston Children’s Hospital and Wheaton College. A complete description of the husbandry and environmental conditions in housing for the fish used in these experiments at Boston Children’s Hospital is available as a collection in protocols.io (https://doi.org/10.17504/protocols.io.mrjc54n). Similar conditions were present at Wheaton College. For all experiments, adult stages were defined by reproductively mature fish >3 months old. Males and females were used together in analyses as both sexes showed comparable changes in growth regulation.

### Photography and measurement of fish, fins, barbels, and fin segments

Fish were anesthetized in MS-222 prior to photography. Whole fish photographs were taken using a Canon EOS 6D camera with a Canon 500D 77 mm close-up lens. Photographs were analyzed using Fiji (Schindelin, et al., 2012) to obtain standard length (STL) determined as the length from the tip of the snout to the posterior end of the caudal peduncle; caudal and anal fin length, determined as the length of the longest fin ray; fin area; and barbel length. For fin segment analysis, fins of anesthetized fish were photographed using a Nikon SMZ800 dissecting microscope and a Spot Insight 2 camera with Spot software. Fin and barbel length were normalized for each fish by dividing by standard length. ANOVA was used to compare mean fin and/or barbel length/ STL among different genotypes of fish. Fin segment lengths were measured using Fiji; a minimum of 20 segments were measured per fish. Segment length data from a minimum of 4 fish were used to calculate mean and SEM.

### Mutagenesis, modifier screen, identification, and genotyping of *schleier*

Homozygous *longfin* (dt2) mutant males were mutagenized with 3.3µM N ethyl-n-nitrosourea for four repeated doses following (Rohner, et al., 2011). Founder mutagenized males were crossed to Tubingen wildtype females. F1 progeny were scored for reduction or increase of heterozygous *longfin* phenotype. An enhancer mutant was identified that led to an increase in all fins and barbs in the heterozygous *lof* background. This mutant was outcrossed and isolated as an independent locus affecting fin growth. The mutant was named *schleier* and given the allele assignment *dmh39*. The mutation underlying the *schleier* mutant phenotype was identified through whole genome sequencing and mapped though identity by descent as previously outlined (Bowen, et al., 2012). The C583Y variant was identified in *slc12a7a* on chromosome 2 and found to be tightly linked to the *schleier* phenotype.

Genotyping the C583Y *slc12a7a/kcc4a* mutation was carried out as follows: DNA was extracted from fin clips using the NaOH method of Meeker *et al*. DNA was denatured at 95°C and snap cooled on ice, then used as template for PCR amplification using PrimeSTAR DNA polymerase (Takara) under manufacturer conditions. Genotyping primers are as follows: F: 5’- GTACAGGTTTTTCTTGATGT-3’, R: 5’-GAAAGGCTCAGGCTCATCCC-3’. PCR products were Sanger sequenced using the reverse primer.

### Creation of *slc12a7a/kcc4a* and *kcnk5b* knockouts

For *slc12a7a*, the online tool ZiFit was used to design guides to target exon 3 of isoform 002 of *D. rerio slc12a7a*. This exon is shared by all 3 major splice isoforms of the gene. The target sequence 5’-GGGGGCTGTCGAGCACGAGG-3’ was used for designing primers TAGGGGGCTGTCGAGCACGAGG and AAACCCTCGTGCTCGACAGCCC which were synthesized, annealed and placed into the BsaI-digested vector pDR274 (Addgene) using the method of Hwang *et al*. *Slc12a7a* guide RNA was synthesized from this vector using the MEGAshortscript T7 Kit (Ambion) as per manufacturer’s instructions.

For *kcnk5b*, the online tool CHOPCHOP was used to design two guides to target exon 1 of *D. rerio kcnk5b*. The two target sequences were 5’-TGCGATATTCCAAATCCTCGAGG-3’ and 5’-ATATCCCTGCCTAAGTAAAGAGG-3’. Guide RNAs (crRNA) for the target sequences and tracrRNA were synthesized by Integrated DNA Technologies. RNAs were resuspended in nuclease-free H_2_0 at 100 uM concentration. The crRNA and tracrRNA were then annealed in manufacturer-provided duplex buffer by heating to 95°C and cooling to room temperature at 20 uM concentration.

Guide RNAs were mixed with Cas9 SmartNuclease eukaryotic mRNA (System Biosciences) such that the final concentration of RNA was approximately 350 ng/uL for each gene (150 ng/ul guide RNA, 300 ng/ul Cas9 RNA). This mixture was injected into single-cell *D. rerio* wild-type (AB) embryos or *schleier/+* embryos, depending on the experiment. After two days, DNA from individual embryos was extracted using the NaOH method of Meeker *et al* and analyzed by PCR and Sanger sequencing. Exon 3 of *slc12a7a* was PCR amplified using the primers F: 5’-GCCGTAAATAAGGTG-3’, R: 5’-CTGTGTGGGAGTCTAGTTG-3’. Exon 1 of *kcnk5b* was PCR amplified using the primers F: 5’-AGAAACTTGGGAGTGTGGAGTG-3’, R: 5’-TTCCCAAAACACAAATGAAACA-3’; this primer pair will detect mutations introduced by either *kcnk5b* guide. PCR products were subcloned into pGEM-T and individual clones were sequenced to determine whether mutations had been introduced at the targeted sequence.

Injected fish (P0) were grown to maturity and used for fin length analyses. *Slc12a7a-* injected P0 fish were subsequently crossed to wild-type AB fish to generate F1 progeny with unique mutations. When DNA analysis was required, anesthetized fish were subjected to fin clip, and DNA extraction and subsequent analysis was carried out on fin tissue as above to identify mutations.

### Construction of Slc12a7a/Kcc4a expression vector

Full length *D. rerio slc12a7a/kcc4a* was PCR amplified from cDNA of wildtype fin and subcloned in pGEM-T (F: 5’-GCACTTTTGTCTCCGTGTCAGC-3’, R: 5’-CATCTTGCTGGAACAGAAACTGAGG-3’). Copy cutter cells (Epicentre) were used for propagation as the *slc12a7a/kcc4a* gene product was toxic to commonly used bacteria when at high copy number. The *slc12a7a/kcc4a* cDNA was subcloned into the *Xenopus* expression vector pXT7 (Addgene). Subsequently, the C583Y mutation was introduced into the *slc12a7a/kcc4a* pXT7 construct using a QuikChange II Kit (Agilent) as per the manufacturer’s protocol.

### Assessment of the K+:Cl− cotransporter function and immunoblot

K+:Cl− cotransport activity of wild-type Kcc4a and its mutant C583Y was assessed using the heterologous expression system of *Xenopus laevis* oocytes following our standard procedures (Adragna, et al., 2015; Melo, et al., 2013; Kahle, et al., 2005). Briefly, mature oocytes were injected with water alone or containing 0.2 µg/µl of Kcc4a cRNA alone or together with its mutant C583Y. Two days after injection, the activity of K+:Cl− cotransport was determined by measuring the Cl--dependent 86Rb+ uptake under hypotonic conditions (∼110 mOsm). All transport assays were performed with groups of 10-15 oocytes pre-incubated for 30 min at room temperature in a Na+ and Cl− free medium (in mM: 50 N-methyl-D-glucamine-gluconate, 10 K+-gluconate, 4.6 Ca2+-gluconate, 1 Mg+-gluconate, and 5 HEPES/Tris, pH 7.4) and then transferred to the uptake medium (in mM: 40 N-methyl-D-glucamine-Cl−, 10 KCl, 1.8 CaCl2, 1 MgCl2, and 5 HEPES/Tris, pH 7.4) containing 0.5 µCi of 86Rb+/ml for 1 h at 32°C. Tracer activity was determined for each oocyte by β-scintillation counting.

For the immunoblot, total extracts of two oocytes equivalents injected with WT or C583Y cRNA were obtained using ice-cold lysis buffer (in mM: 50 Tris-HCl (pH 7.5), 1 EGTA, 1 EDTA, 50 NaF, 5 Na4O7P2, 1 Na3VO4, 1% (w/v) Nonidet P40, 0.27 sucrose, 0.1% 2-mercaptoethanol) supplemented with a protease inhibitor cocktail (1 tablet per 50 ml, Sigma). Total KCC4 antibody used was previously described (Adragna, et al., 2015; Melo, et al., 2013; Kahle, et al., 2005), while β-actin and secondary antibody coupled to horseradish peroxidase used were obtained from Santa Cruz.

### Analysis of vascular flow

To compare flow rate and diameter of vessels, three wildtype and three *schleier* fish were analyzed (mass = 0.48 +/-0.02 g; STL = 26 +/-0.5 mm; +/- = SEM). Each fish was lightly anesthetized by submersion in a dilute concentration of 0.04% MS-222; 0.01 ml of 0.005 mg/ml pancuronium bromide (Sigma) was then injected intraperitoneally for immobilization (Arnaout, et al., 2007), as even minor movements caused video artifacts. Fish were then transferred to an MS-222 soaked sponge with a small cutout; this set-up stabilized the fish while allowing for the maintenance of anesthesia and a moist environment. Each fish was kept in this setup for a maximum of ten minutes. Heart rate was recorded before and after data collection to verify that the experimental procedure did not alter heart rate. Throughout the experiment, diluted anesthetic was repeatedly introduced over the gills to ensure the fish remained anesthetized and well-ventilated. For data collection, the fish was moved to the stage of a Nikon Eclipse E400 microscope which was fitted with a HiSpec Lite camera to record at 200 fps with a resolution of 704 × 528 pixels. The fish was filmed at the posterior-most tip of the caudal fin under 200X magnification. Fish were euthanized by MS-222 overdose immediately following video recording.

From the video data, blood vessel diameter and red blood cell velocity were measured using ImageJ software (Rueden, et al., 2017). Blood vessel diameter was measured near the tip of the caudal fin for the major artery running through the second-most dorsal fin ray and from the paired vein directly above it. The diameter of each vessel was measured in at least five positions and data were pooled per individual fish. From these measurements, a ratio of vein:artery diameter for each fish was calculated and compared between wildtype and *schleier* fish using an ANOVA. Red blood cell velocity was similarly measured at the tip of the caudal fin artery and vein. For each vessel, a minimum of ten red blood cells were tracked for 10-50 frames (50-250ms) each by using the MTrackJ plugin https://imagescience.org/meijering/software/mtrackj/. Data were pooled within individual fish for each velocity location. As was performed for the diameter analysis, a ratio of vein:artery red blood cell velocity was calculated per fish and compared between wildtype and *schleier* fish using an ANOVA.

### Immunofluorescence and *in situ* hybridization

Caudal fins were amputated using a razor blade from adult fish anesthetized with 0.04% MS-222. Fins were fixed for one hour at room temperature and then at 4°C overnight in a solution of 4% paraformaldehyde/ PBS, pH 7.4. After overnight fixation, 0.5 M EDTA solution was added to the sample for a final concentration of 0.25 M EDTA and incubated for an additional 24 hours at 4°C. Fins were rinsed several times in PBS and moved gradually through a sucrose/PBS solutions of increasing sucrose concentration (15% × 2 hours, 20% × 2 hours, 30% overnight) while kept at 4°C. The following day, fins were transferred to cryo-embedding medium (Tissue-Tek O.C.T compound) in cryomolds and allowed to equilibrate for at least 1 hour at room temperature. Samples were then flash frozen in isopentane chilled in a dry ice-isopentane bath, and transferred to −80°C. For cryosectioning, blocks were allowed to warm to –20°C, then sectioned into 10 uM transverse sections using a Microm HM 505N microtome cryostat. Sections were collected onto Fisherbrand ColorFrost Plus slides, dried for 1 hour at 37°C, dried overnight at room temperature, and then stored at −80°C for up to 3 months.

#### Immunofluorescence

Slides were thawed at room temperature. After drying slightly, an ImmunoPen (Sigma) was used to isolate individual sections. Sections were incubated in blocking solution (5% goat serum in PBS) in a humidified chamber for one hour at room temperature. After block was removed, primary antibody (KCC4/SLC12A7 rabbit polyclonal antibody, Novus Biologicals NBP1-49583) was added at 1:100 dilution in PBST (1% goat serum, 0.3% Tween in PBS). Slides were incubated in primary antibody in a humidified chamber overnight at 4°C. Slides were washed gently in PBST and then incubated in secondary antibody at 1:200 dilution (Goat anti-Rabbit IgG (H+L) Secondary Antibody, Alexa Fluor® 555 conjugate, ThermoFisher) in PBST for 1 hour at room temperature in a dark humidified chamber. Slides were washed gently in PBST, mounted in Prolong Diamond antifade mount with DAPI (ThermoFisher) and glass coverslips, and dried overnight at room temperature in the dark. Slides were imaged using a Nikon Eclipse E400 microscope equipped for fluorescence detection and photographed with a Spot Insight 2 Camera.

#### *In situ* hybridization

Slides were thawed at room temperature, then post-fixed in 4% paraformaldehyde/ PBS, pH 9.5 for 1 hour at room temperature. Following fixation, slides were rinsed thoroughly in PBS in Coplin jars, dehydrated in an ethanol series (70%, 95%, 100%) and allowed dry at room temperature before permeabilization with 1% Triton X-100 in PBS (20 minutes, room temperature). Slides were rinsed again in PBS and dehydrated in ethanol as above. Slide edges were sealed with an ImmunoPen.

Hybridization procedures were modified from (Smith, et al., 2008). Probes were diluted 1:150 in hybridization buffer (1X salt [0.2 M NaCl, 10 mM Tris HCl, 5 mM NaH_2_PO_4_, 5 mM Na_2_HPO_4_, 1 mM Tris base, 5 mM EDTA], 50% deionized formamide, 10% dextran sulphate, 0.05 mg/ml yeast tRNA, and 1× Denhardts). Probes were heated for 10 minutes at 70°C, then placed on ice briefly before applying to slides. Slides with probe were incubated flat in RNAse-free containers humidified with 1X salt/50% formamide at 55°C for 48 hours. After incubation, slides were washed in solution A (1× SSC, 50% formamide, and 0.1% Tween 20) in slide mailers three times at 55°C (15 minutes, 30 minutes, 30 minutes), then washed twice in MABT (100 mM maleic acid, 150 mM NaCl, 0.1% Tween-20, pH 7.5) at room temperature for 30 minutes per wash). Slides were placed flat in containers humidified with dH_2_0 and incubated in a small volume of blocking solution (20% heat-inactivated sheep serum in MABT) overnight at 4°C. Block solution was removed and anti-digoxigenin-AP, Fab fragments (Sigma) diluted 1:2000 in fresh blocking solution was added. Slides were incubated in antibody for a minimum of 24 hours at 4°C. Slides were then washed at room temperature in slide mailers as follows: 4 washes of 20 minutes each in MABT, followed by 2 washes of 10 minutes each in NTMT staining buffer (100 mM NaCl, 100 mM Tris HCL pH 9.5, 50 mM MgCl2, and 0.1% Tween 20). Slides were placed flat in containers humidified with dH_2_0 and incubated in BM Purple (Sigma) in the dark at room temperature for 48 hours. After development of signal, slides were washed in PBS-0.1% Triton and fixed in 4% PFA/PBS overnight at 4°C. Slides were washed in ddH_2_0, allowed to dry briefly, and mounted in CC mount (Sigma) under glass coverslips. After overnight drying, slides were photographed using a Nikon Eclipse E400 microscope with a Spot Insight 2 Camera.

#### *In situ* probe cloning and synthesis

Primer3 was used to design PCR primers in exons 21 and 26 of the *slc12a7a/kcc4a gene* (F: 5’- GATGGAACAACGCTCACAGA-3’; R: 5’-GAGGACCTCCAGAAACTCCA-3’). These primers were used to amplify a 517 base pair PCR product from caudal fin cDNA, which was then subcloned into the pGEM-T vector using TA cloning. For probe generation, 10 ug of purified plasmid was digested with SacI or SacII restriction enzyme, phenol-chloroform extracted, precipitated with sodium acetate/ethanol, and resuspended in nuclease-free H_2_0. 1 ug purified digested plasmid was used as template for SP6 or T7 transcription using the DIG-RNA labeling kit (Roche). Following probe transcription, DNA template was removed using Turbo DNAse kit (In Vitrogen). Probe was precipitated by incubation in 0.1 M LiCl, 75% ethanol overnight at –80°C, followed by centrifugation and wash in 70% ethanol. After drying slightly, probe pellet was resuspended in 50% nuclease-free H_2_0 / 50% formamide and stored at –80°C. An aliquot of probe was analyzed using agarose electrophoresis to verify probes were expected sizes and not degraded.

## References

Adams, D.S., Uzel, S.G., Akagi, J., Wlodkowic, D., Andreeva, V., Yelick, P.C., Devitt-Lee, A., Pare, J.F., and Levin, M. (2016). Bioelectric signalling via potassium channels: a mechanism for craniofacial dysmorphogenesis in KCNJ2-associated Andersen-Tawil Syndrome. J Physiol 594, 3245–70.

Adragna, N.C., Ravilla, N.B., Lauf, P.K., Begum, G., Khanna, A.R., Sun, D., and Kahle, K.T. (2015). Regulated phosphorylation of the K-Cl cotransporter KCC3 is a molecular switch of intracellular potassium content and cell volume homeostasis. Front Cell Neurosci 9, 255.

Arnaout, R., Ferrer, T., Huisken, J., Spitzer, K., Stainier, D.Y., Tristani-Firouzi, M., and Chi, N.C. (2007). Zebrafish model for human long QT syndrome. Proc Natl Acad Sci U S A 104, 11316–21.

Bagriantsev, S.N., Peyronnet, R., Clark, K.A., Honore, E., and Minor, D.L., Jr. (2011). Multiple modalities converge on a common gate to control K2P channel function. EMBO J 30, 3594–606.

Barriere, H., Belfodil, R., Rubera, I., Tauc, M., Lesage, F., Poujeol, C., Guy, N., Barhanin, J., and Poujeol, P. (2003). Role of TASK2 potassium channels regarding volume regulation in primary cultures of mouse proximal tubules. J Gen Physiol 122, 177–90.

Bergeron, M.J., Boggavarapu, R., Meury, M., Ucurum, Z., Caron, L., Isenring, P., Hediger, M.A., and Fotiadis, D. (2011). Frog oocytes to unveil the structure and supramolecular organization of human transport proteins. PLoS One 6, e21901.

Boettger, T., Hubner, C.A., Maier, H., Rust, M.B., Beck, F.X., and Jentsch, T.J. (2002). Deafness and renal tubular acidosis in mice lacking the K-Cl co-transporter Kcc4. Nature 416, 874–8.

Bowen, M.E., Henke, K., Siegfried, K.R., Warman, M.L., and Harris, M.P. (2012). Efficient mapping and cloning of mutations in zebrafish by low-coverage whole-genome sequencing. Genetics 190, 1017–24.

Brown, T.C., Murtha, T.D., Rubinstein, J.C., Korah, R., and Carling, T. (2018). SLC12A7 alters adrenocortical carcinoma cell adhesion properties to promote an aggressive invasive behavior. Cell Commun Signal 16, 27.

Chen, Y.F., Chou, C.Y., Wilkins, R.J., Ellory, J.C., Mount, D.B., and Shen, M.R. (2009). Motor protein-dependent membrane trafficking of KCl cotransporter-4 is important for cancer cell invasion. Cancer Res 69, 8585–93.

Daane, J.M., Lanni, J., Rothenberg, I., Seebohm, G., Higdon, C.W., Johnson, S.L., and Harris, M.P. (2018). Bioelectric-calcineurin signaling module regulates allometric growth and size of the zebrafish fin. Sci Rep 8, 10391.

Dahal, G.R., Pradhan, S.J., and Bates, E.A. (2017). Inwardly rectifying potassium channels influence Drosophila wing morphogenesis by regulating Dpp release. Development 144, 2771–2783.

Dahal, G.R., Rawson, J., Gassaway, B., Kwok, B., Tong, Y., Ptacek, L.J., and Bates, E. (2012). An inwardly rectifying K+ channel is required for patterning. Development 139, 3653–64.

Donaldson, M.R., Yoon, G., Fu, Y.H., and Ptacek, L.J. (2004). Andersen-Tawil syndrome: a model of clinical variability, pleiotropy, and genetic heterogeneity. Ann Med 36 Suppl 1, 92–7.

Felix, R. (2000). Channelopathies: ion channel defects linked to heritable clinical disorders. J Med Genet 37, 729–40.

Ferre, F., and Clote, P. (2005). DiANNA: a web server for disulfide connectivity prediction. Nucleic Acids Res 33, W230–2.

Gokhale, R.H., and Shingleton, A.W. (2015). Size control: the developmental physiology of body and organ size regulation. Wiley Interdiscip Rev Dev Biol 4, 335–56.

Hoptak-Solga, A.D., Klein, K.A., DeRosa, A.M., White, T.W., and Iovine, M.K. (2007). Zebrafish short fin mutations in connexin43 lead to aberrant gap junctional intercellular communication. FEBS Lett 581, 3297–302.

Hsu, Y.M., Chou, C.Y., Chen, H.H., Lee, W.Y., Chen, Y.F., Lin, P.W., Alper, S.L., Ellory, J.C., and Shen, M.R. (2007). IGF-1 upregulates electroneutral K-Cl cotransporter KCC3 and KCC4 which are differentially required for breast cancer cell proliferation and invasiveness. J Cell Physiol 210, 626–36.

Huxley, J. (1932). Problems in Relative Growth The Dial Press)

Huxley, J., and Teissier, G. (1936). Terminology of Relative Growth. Nature 137, 780–781.

Inaba, M., Yamanaka, H., and Kondo, S. (2012). Pigment pattern formation by contact-dependent depolarization. Science 335, 677.

Iovine, M.K., Higgins, E.P., Hindes, A., Coblitz, B., and Johnson, S.L. (2005). Mutations in connexin43 (GJA1) perturb bone growth in zebrafish fins. Dev Biol 278, 208–19.

Irion, U., Frohnhofer, H.G., Krauss, J., Colak Champollion, T., Maischein, H.M., Geiger-Rudolph, S., Weiler, C., and Nusslein-Volhard, C. (2014). Gap junctions composed of connexins 41.8 and 39.4 are essential for colour pattern formation in zebrafish. Elife 3, e05125.

Iwashita, M., Watanabe, M., Ishii, M., Chen, T., Johnson, S.L., Kurachi, Y., Okada, N., and Kondo, S. (2006). Pigment pattern in jaguar/obelix zebrafish is caused by a Kir7.1 mutation: implications for the regulation of melanosome movement. PLoS Genet 2, e197.

Kahle, K.T., Rinehart, J., de Los Heros, P., Louvi, A., Meade, P., Vazquez, N., Hebert, S.C., Gamba, G., Gimenez, I., and Lifton, R.P. (2005). WNK3 modulates transport of Cl− in and out of cells: implications for control of cell volume and neuronal excitability. Proc Natl Acad Sci U S A 102, 16783–8.

Lang, F., and Stournaras, C. (2014). Ion channels in cancer: future perspectives and clinical potential. Philos Trans R Soc Lond B Biol Sci 369, 20130108.

Lawson, N.D., and Weinstein, B.M. (2002). In vivo imaging of embryonic vascular development using transgenic zebrafish. Dev Biol 248, 307–18.

Levin, M. (2014). Endogenous bioelectrical networks store non-genetic patterning information during development and regeneration. J Physiol 592, 2295–305.

Levin, M., Pezzulo, G., and Finkelstein, J.M. (2017). Endogenous Bioelectric Signaling Networks: Exploiting Voltage Gradients for Control of Growth and Form. Annu Rev Biomed Eng 19, 353–387.

Litan, A., and Langhans, S.A. (2015). Cancer as a channelopathy: ion channels and pumps in tumor development and progression. Front Cell Neurosci 9, 86.

Lobikin, M., Chernet, B., Lobo, D., and Levin, M. (2012). Resting potential, oncogene-induced tumorigenesis, and metastasis: the bioelectric basis of cancer in vivo. Phys Biol 9, 065002.

Marcoux, A.A., Garneau, A.P., Frenette-Cotton, R., Slimani, S., Mac-Way, F., and Isenring, P. (2017). Molecular features and physiological roles of K(+)-Cl(-) cotransporter 4 (KCC4). Biochim Biophys Acta Gen Subj 1861, 3154–3166.

McClenaghan, C., Schewe, M., Aryal, P., Carpenter, E.P., Baukrowitz, T., and Tucker, S.J. (2016). Polymodal activation of the TREK-2 K2P channel produces structurally distinct open states. J Gen Physiol 147, 497–505.

McLaughlin, K.A., and Levin, M. (2018). Bioelectric signaling in regeneration: Mechanisms of ionic controls of growth and form. Dev Biol 433, 177–189.

Melo, Z., de los Heros, P., Cruz-Rangel, S., Vazquez, N., Bobadilla, N.A., Pasantes-Morales, H., Alessi, D.R., Mercado, A., and Gamba, G. (2013). N-terminal serine dephosphorylation is required for KCC3 cotransporter full activation by cell swelling. J Biol Chem 288, 31468–76.

Misu, A., Yamanaka, H., Aramaki, T., Kondo, S., Skerrett, I.M., Iovine, M.K., and Watanabe, M. (2016). Two Different Functions of Connexin43 Confer Two Different Bone Phenotypes in Zebrafish. J Biol Chem 291, 12601–11.

Nusslein-Volhard, C., and Dahm, R. (2002). Zebrafish: a practical approach (Oxford Oxford University Press)

Othmer, H.G., and Pate, E. (1980). Scale-invariance in reaction-diffusion models of spatial pattern formation. Proc Natl Acad Sci U S A 77, 4180–4.

Perathoner, S., Daane, J.M., Henrion, U., Seebohm, G., Higdon, C.W., Johnson, S.L., Nusslein-Volhard, C., and Harris, M.P. (2014). Bioelectric signaling regulates size in zebrafish fins. PLoS Genet 10, e1004080.

Prevarskaya, N., Skryma, R., and Shuba, Y. (2010). Ion channels and the hallmarks of cancer. Trends Mol Med 16, 107–21.

Rohner, N., Perathoner, S., Frohnhofer, H.G., and Harris, M.P. (2011). Enhancing the efficiency of N-ethyl-N-nitrosourea-induced mutagenesis in the zebrafish. Zebrafish 8, 119–23.

Rueden, C.T., Schindelin, J., Hiner, M.C., DeZonia, B.E., Walter, A.E., Arena, E.T., and Eliceiri, K.W. (2017). ImageJ2: ImageJ for the next generation of scientific image data. BMC Bioinformatics 18, 529.

Schindelin, J., Arganda-Carreras, I., Frise, E., Kaynig, V., Longair, M., Pietzsch, T., Preibisch, S., Rueden, C., Saalfeld, S., Schmid, B., et al. (2012). Fiji: an open-source platform for biological-image analysis. Nat Methods 9, 676–82.

Shen, M.R., Chou, C.Y., Hsu, K.F., Liu, H.S., Dunham, P.B., Holtzman, E.J., and Ellory, J.C. (2001). The KCl cotransporter isoform KCC3 can play an important role in cell growth regulation. Proc Natl Acad Sci U S A 98, 14714–9.

Simard, C.F., Bergeron, M.J., Frenette-Cotton, R., Carpentier, G.A., Pelchat, M.E., Caron, L., and Isenring, P. (2007). Homooligomeric and heterooligomeric associations between K+-Cl−cotransporter isoforms and between K+-Cl− and Na+-K+-Cl− cotransporters. J Biol Chem 282, 18083–93.

Smith, A., Zhang, J., Guay, D., Quint, E., Johnson, A., and Akimenko, M.A. (2008). Gene expression analysis on sections of zebrafish regenerating fins reveals limitations in the whole-mount in situ hybridization method. Dev Dyn 237, 417–25.

Umulis, D.M., and Othmer, H.G. (2013). Mechanisms of scaling in pattern formation. Development 140, 4830–43.

Vollmer, J., Casares, F., and Iber, D. (2017). Growth and size control during development. Open Biol 7.

Wang, Z. (2004). Roles of K+ channels in regulating tumour cell proliferation and apoptosis. Pflugers Arch 448, 274–86.

Warth, R., Barriere, H., Meneton, P., Bloch, M., Thomas, J., Tauc, M., Heitzmann, D., Romeo, E., Verrey, F., Mengual, R., et al. (2004). Proximal renal tubular acidosis in TASK2 K+ channel-deficient mice reveals a mechanism for stabilizing bicarbonate transport. Proc Natl Acad Sci U S A 101, 8215–20.

Watanabe, M., Iwashita, M., Ishii, M., Kurachi, Y., Kawakami, A., Kondo, S., and Okada, N. (2006). Spot pattern of leopard Danio is caused by mutation in the zebrafish connexin41.8 gene. EMBO Rep 7, 893–7.

Watt, K.I., Harvey, K.F., and Gregorevic, P. (2017). Regulation of Tissue Growth by the Mammalian Hippo Signaling Pathway. Front Physiol 8, 942.

Yakar, S., Werner, H., and Rosen, C.J. (2018). Insulin-like growth factors: actions on the skeleton. J Mol Endocrinol 61, T115–T137.

Zhou, C., Feng, X., Yuan, F., Ji, J., Shi, M., Yu, Y., Zhu, Z., and Zhang, J. (2018). Difference of molecular alterations in HER2-positive and HER2-negative gastric cancers by whole-genome sequencing analysis. Cancer Manag Res 10, 3945–3954.

