## Supplementary figures and captions for "Integrated K^+^ channel and K^+^-Cl^−^ cotransporter functions regulate fin proportionality in zebrafish"

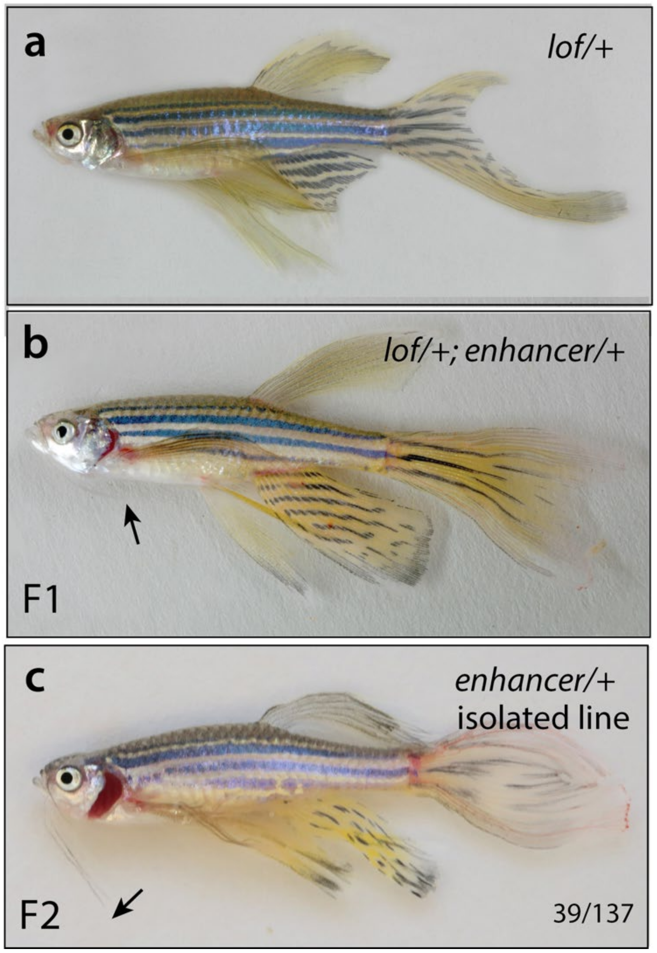


**Supplementary Figure 1. Identification of an enhancer of longfin and additive effects of fin growth**. Mutagenized wild-type founder fish were outcrossed to *lof* homozygotes and 1081 F1 progeny were isolated. 16 fish were isolated with altered *longfin* phenotypes. **a**) *lof* heterozygous phenotype. **b**) Founder fish identified with enhancer mutation in *trans* to *lof* with increased size of all fins. Barbules were also found to be longer (arrows), a trait not seen in *lof* homozygotes. **c)** Isolated strain of enhancer line, named *schleier/dmh39,* segregating independently from *lof*.

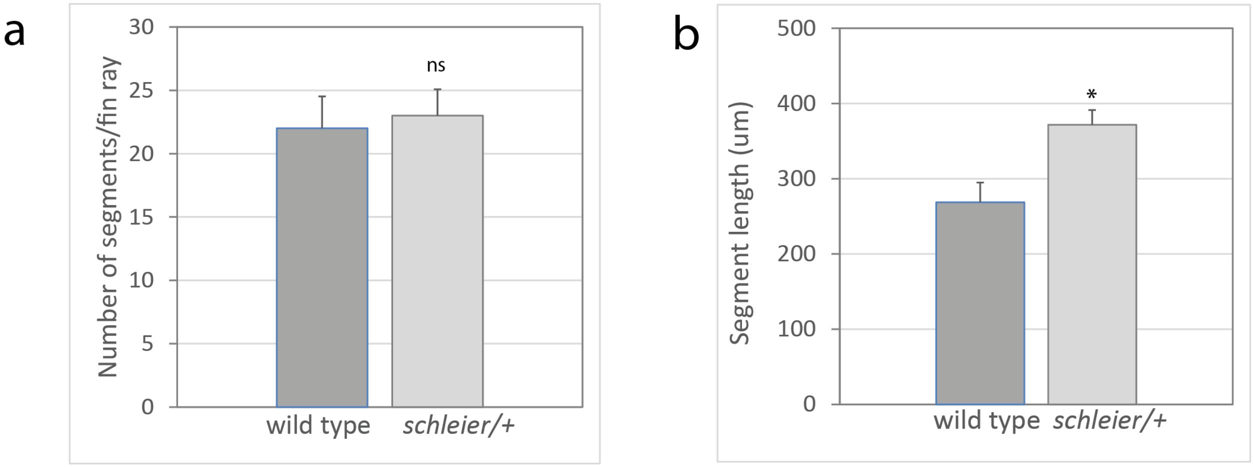


**Supplementary Figure 2. Segment number and length in wild-type and *schleier* heterozygous fish.** Caudal fin ray segments of adult wild-type or *schleier* heterozygous fish were counted and measured across the proximodistal axis. **a)** Number of segments per fin ray; mean +/- SEM is shown. n.s., not significant; n=3 fish per group. **b**) Segment length per genotype; mean +/- SEM is shown. *, $p<0.05$ calculated using a student's T test; n=3 fish per group, ≥17 segments measured per fish.

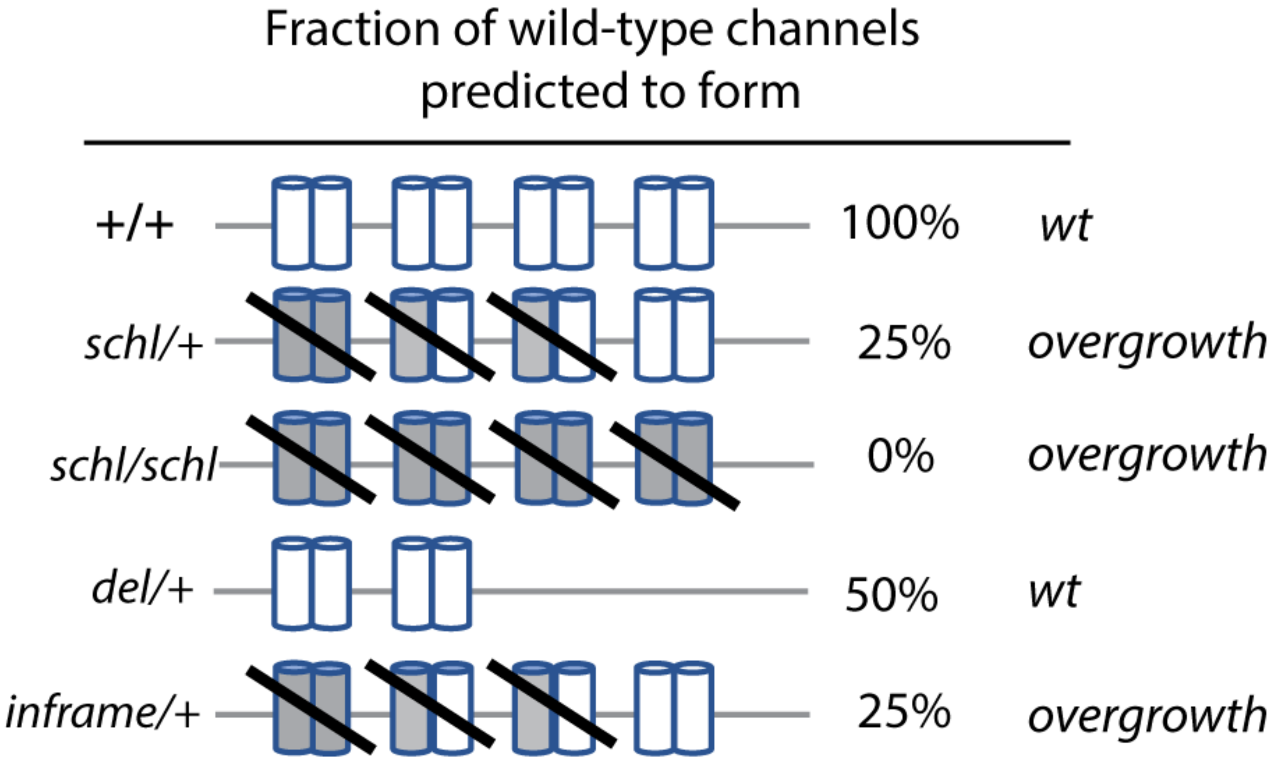


**Supplementary file 3.** Genetic regulation of overgrowth phenotype by Kcc4a allelic variations through reduction of wild-type functioning Kcc4a. Percentage indicates predicted amount of functional Kcc4a protein present in each genotype; overgrowth and wt (wild-type) refer to expected fin phenotype.

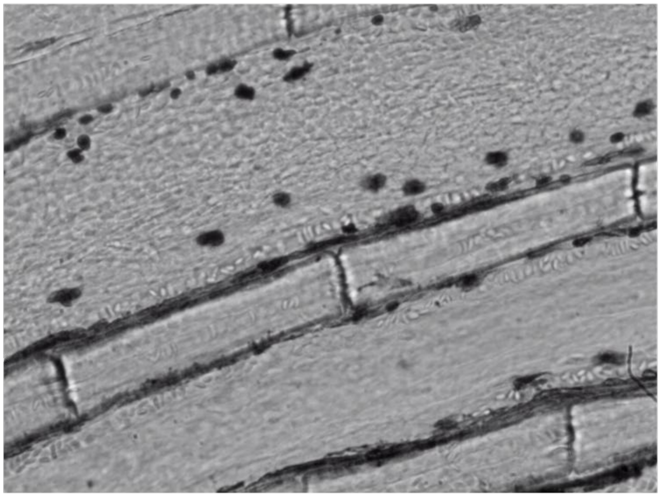

**Supplementary Movie 1. Velocity and volume of blood flow is similar in wild-type fin arteries and veins.** Video shows recording of wild-type caudal fin under 200X magnification; anterior-posterior orientation is left to right; dorsal-ventral orientation is top to bottom. Blood flow in caudal fin artery proceeds from left (fin base) to right (fin tip). Video playback speed = 30 frames per second (fps); recording speed = 200 fps.

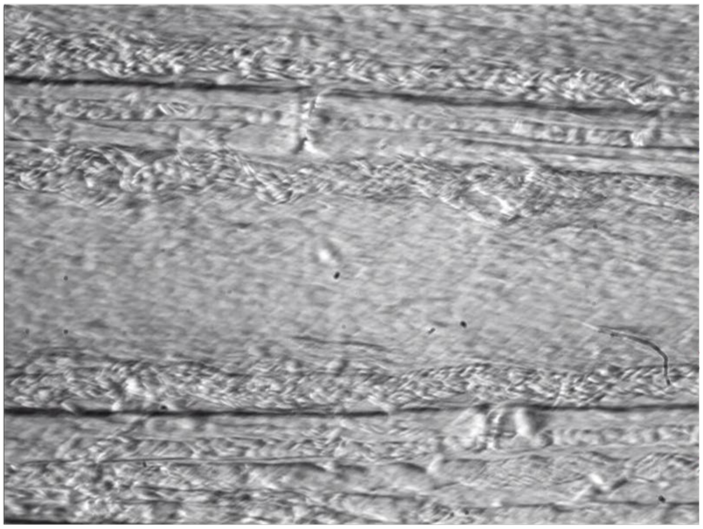

**Supplementary Movie 2. Velocity and volume of blood flow are altered in *schleier* fin arteries and veins.** Video shows recording of *schleier*/+ caudal fin under 200X magnification; anterior-posterior orientation is left to right; dorsal-ventral orientation is top to bottom. Blood flow in caudal fin artery proceeds from left (fin base) to right (fin tip). Note decreased velocity and increased volume of blood flow, particularly in veins. Video playback speed = 30 frames per second (fps); recording speed = 200 fps.
